# General Patterns in the Temperature-Dependence of Heterotrophic Microbial Interactions

**DOI:** 10.1101/2025.04.18.649480

**Authors:** Quqiming Duan, Will Harcombe, Van Savage, Michael Mustri, Thomas P. Smith, Samraat Pawar

## Abstract

Despite its global importance, our ability to predict the impacts of temperature change on the community dynamics of heterotrophic microbes remains limited. Here, we develop a metabolic trait-based mathematical framework to predict the temperature dependence of pairwise interactions among heterotrophic microbial consumers, accounting for their resource environment and community composition. Applying this framework leads to two general predictions. First, microbial species interactions are typically more thermally sensitive than the underlying metabolic traits. Second, temperature systematically reshapes intra- and interspecific interactions: their variance peaks at intermediate temperatures, while mean interaction strengths increase with warming more rapidly than interspecific interaction strengths. We show that these features of temperature-dependent interactions have far-reaching implications for the temperature responses of community coexistence, diversity, and stability. Our framework provides a mechanistic foundation for predicting how temperature affects the dynamics of heterotrophic microbial communities across diverse biological and environmental contexts.

**Significance:** Communities of heterotrophic microbes, including bacteria, archaea, protists, and fungi, play a fundamental role in human health, bioprocessing, and global biogeochemical cycles. Predicting their responses to environmental change is a major challenge, with a key missing link being the effects of temperature on species interactions that ultimately shape community dynamics. By integrating metabolic constraints into consumer-resource theory, we derive general predictions about how interactions among heterotrophic microbial species respond to temperature changes. We show that interactions are more sensitive to warming than individual metabolic traits and undergo systematic temperature-dependent changes that alter community-level coexistence, diversity, and stability. These results hold across diverse environmental contexts and heterotrophic systems, providing a foundation for predicting how microbial community dynamics respond to environmental temperature, from biotechnological applications to natural ecosystems.

## 1 Introduction

One of nature’s most profound facts is that no organism exists in isolation. This is especially true for microbes, which in the real world typically exist in complex communities structured by interactions between trillions of cells, thousands of strains, and hundreds of species (Thompson et al. 2017, Machado et al. 2021, The Human Microbiome Project Consortium et al. 2012). Microbial species interactions are particularly crucial because Bacteria, Archaea, Fungi, and Protists collectively dominate heterotrophic biomass in practically all ecosystems (Bar-On et al. 2018), driving the recycling of matter and energy (Philippot et al. 2013, Handley 2019, Malik et al. 2020). In all of this, prokaryotes, and in particular bacteria, play a pivotal role due to their high biomass abundance relative to other microbial groups (Bar-On et al. 2018, Rivett & Bell 2018).

Heterotrophic microbes interact with each other through resource use by modifying their shared resource environment. These interactions mainly include direct competition for shared resources, cross-feeding on each other’s metabolic byproducts (e.g., facilitation, mutualism), and higher-order effects arising from changes in the shared resource environment. At the same time, key metabolic traits—resource consumption and allocation, growth, and respiration rates—all respond unimodally to temperature (Fig. 1; Dell et al. 2011, Corkrey et al. 2012, Smith et al. 2019, Arnoldi et al. 2025). Therefore, environmental temperature is a ubiquitous and fundamental driver of interaction structure, that is, the type (e.g., competition versus cooperation) and strength of microbial species interactions (Bestion et al. 2018, Kaur & Dutta 2020, García et al. 2023, Clegg & Pawar 2024). Temperature-driven changes in the structure of intra- and interspecific interactions can, in turn, affect microbial community dynamics, from functioning to coexistence and stability (García et al. 2023, Pold et al. 2020, Clegg & Pawar 2024). Therefore, predicting the temperature dependence of microbial interactions is necessary for understanding spatio-temporal variation in microbial communities in both natural and artificial (e.g., bioreactor) environments.

**Figure 1.**
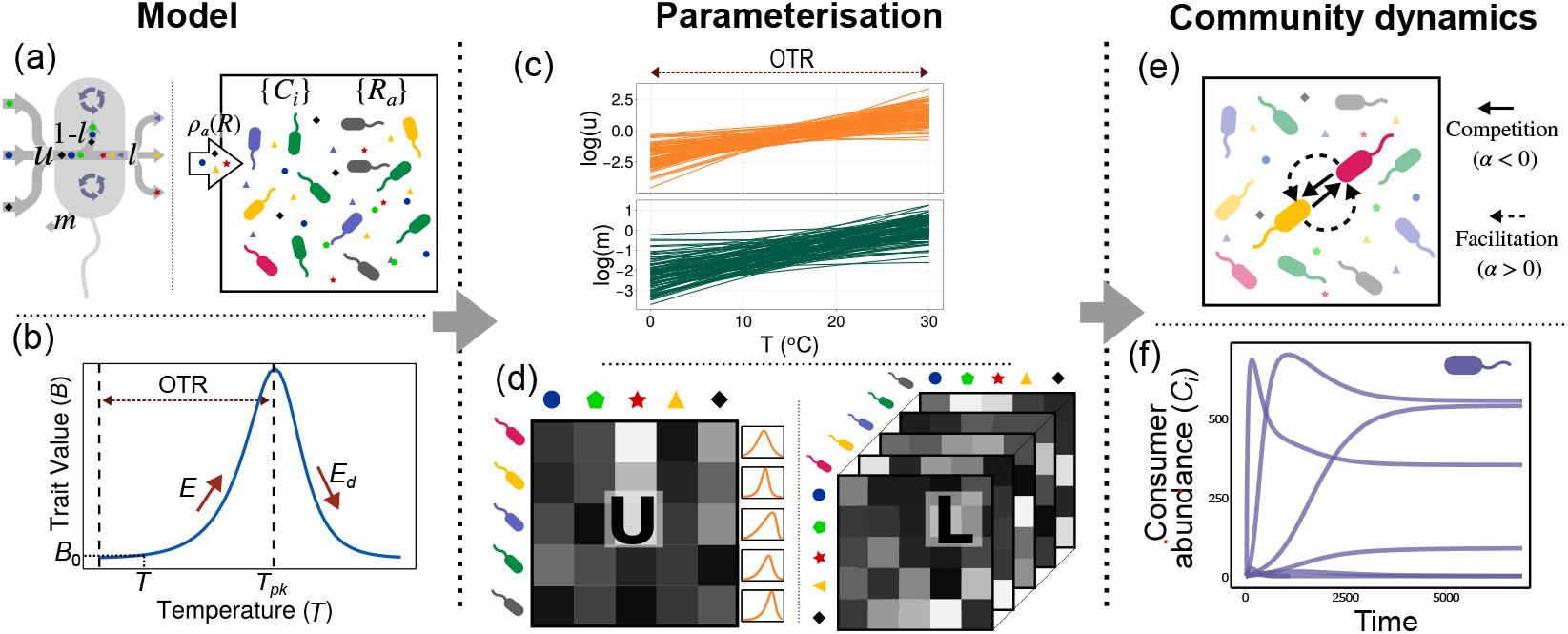
Model schematic and analysis framework. The MiCRM **(a)** and form of the temperature performance curve (TPC) of its two metabolic traits — uptake *u* and respiration rate *m* (Eqns. 3-4) **(b). (c)** Empirically constrained variation in the TPCs of metabolic traits; TPCs within the operational temperature range (OTR) are shown in log-scale. The TPCs determine uptake rates in the matrix **U** and therefore effective metabolite leakage through the tensor **L (d)**. This, in turn, drives the effective interactions between consumer pairs **(e)** and their population dynamics **(f)**.

Our ability to derive or predict the temperature-dependence of interactions between heterotrophic microbial species pairs remains limited. The handful of studies that have sought to tackle this challenge have either relied on empirical inferences of pairwise species interactions based on co-occurrences (Cazelles et al. 2016, Barberán et al. 2012) or made phenomenological assumptions based on the temperature dependence of single-species’ growth and respiration rates (Lax et al. 2020, Clegg & Pawar 2024). The empirical approach has limited generalizability because several confounding factors can lead to species co-occurrence, compounded by the difficulty of obtaining sufficient temperature-specific snapshots of community composition for inference (Blanchet et al. 2020). The second, metabolic-traits-based phenomenological approach can at best predict the qualitative nature of the temperature dependence of pairwise interactions, because it does not consider the consumer-resource dynamics through which these interactions emerge (MacArthur 1969, Marsland et al. 2019, Goldford et al. 2018). On the other hand, significant theoretical progress has been made towards predicting resource-mediated, temperature-dependent interactions between multicellular eukaryotic species from their underlying metabolic traits (Vasseur & McCann 2005, Dell et al. 2014, Gilbert et al. 2014), but these cannot be applied to microbial interactions, which are subject to very different metabolic and biophysical constraints (Stark et al. 2024, DeLong et al. 2010, Kempes et al. 2017, Bestion et al. 2018).

In this study, we develop an approach to derive pairwise microbial species interactions arising from temperature-dependent resource utilization dynamics through a minimal set of empirically feasible metabolic trait measurements. Using this, we reveal general patterns in the temperature dependence of interactions expected to emerge from underlying metabolic traits and show that these patterns have wide-ranging consequences for microbial community dynamics.

## 2 Methods

To derive the temperature-dependence of resource-mediated species interactions in the context of their community composition and resource environment, we incorporate thermal performance curves (TPCs; Fig. 1) of species-level traits into a general consumer-resource model. We then calculate the effective Lotka-Volterra approximation for the system to obtain the temperature dependence of pairwise interactions arising from the underlying metabolically constrained consumer-resource dynamics.

### 2.1 The Consumer-Resource Model

We use the general Microbial Consumer-Resource Model (MiCRM) (Goldford et al. 2018, Marsland et al. 2020*b*, Lechón-Alonso et al. 2021) (Fig. 1):

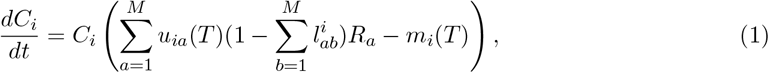

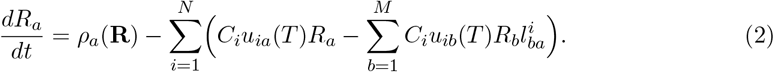

Here, for a community of *N* species (or “strains”) and *M* resource types, *C*_*i*_ (*i* = 1, …, *N* ) is the biomass abundance of the *i*^*th*^ consumer species, and *R*_*a*_ (*a* = 1, …, *M* ) is the resource abundance of the *a*^*th*^ resource type. The function *ρ*_*a*_(*R*_*a*_) is the resource supply, which can take several forms (constant input only, chemostat, leaching, etc.; section S7). The *i*^*th*^ population’s growth is determined by its two temperature (*T* )-dependent traits: resource uptake *u*_*i*_(*T* ) rate, and respiratory loss rate *m*_*i*_(*T* ) to growth and maintenance, assuming that the contribution of cell death to the biomass pool is negligible. The MiCRM is general in that it provides a unified, minimal framework that flexibly captures variation in resource competition and cross-feeding, as well as in energy-use traits and resource-supply types. This is important because, as we show in sections S7-S10, our approach can be applied to the resulting predictions, which hold across a wide range of thermal metabolic constraints, consumer-resource types, and environmental scenarios.

We model the external resource supply *ρ*_*a*_ as a temperature-independent linear function (Cui et al. 2024) because, in most microbial communities, external resource inputs operate on much slower timescales than the rapid metabolic responses of the microbes (e.g., detritus deposition in soils). This implicit separation of time scales in temperature dependence allows us to focus on how consumer uptake (*u*_*i*_(*T* )) and respiration (*m*_*i*_(*T* )) rates govern interaction and community dynamics over the timescales of standard microbial lifespans.

Total resource uptake *u*_*i*_ is determined by its vector of resource preferences (of *u*_*ia*_’s; Fig. 1). A fraction of this is “leaked” and returned to the resource pool, either due to substrate-specific inefficiency 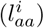 or transformation into metabolic by-products 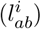, determining the effective rate at which consumed resources are returned to the resource pool. We assume that leakage fractions do not vary with metabolic rate or temperature and are therefore treated as temperature-independent. However, the total effective resource leakage rate remains temperature-dependent because it is proportional to uptake, i.e., 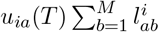. The **U** matrix, consisting of the *N* species’ uptake rate vectors (of *M* resource types) thus represents species-level variation in core metabolic capabilities (Fig. 1d).

The MiCRM treats resources as substitutable (Cui et al. 2024), meaning they satisfy equivalent metabolic and stoichiometric needs such that a species’ growth rate depends on the total resource content harvested rather than on a single limiting substrate or nutrient. While growth can be strictly rate-limited by a single resource, as described by Liebig’s Law of the minimum (De Baar 1994, Gibbs et al. 2022, Klausmeier et al. 2004), our assumption provides a reasonable approximation where growth is limited by a specific group or family of resources (Butler & O’Dwyer 2018).

#### Adding Thermal Metabolic Constraints

We use the modified Sharpe-Schoolfield equation (Schoolfield et al. 1981, García et al. 2023, Smith et al. 2021) to model the TPCs of the MiCRM’s two metabolic parameters (Fig. 1):

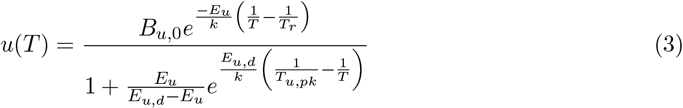

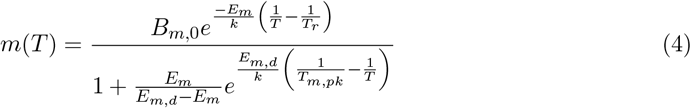

This model assumes that enzyme-catalyzed physiological rates follow Arrhenius kinetics over the low-to-medium operational temperature range (Fig. 1b), with activation energy *E* and reversible thermal inactivation of the key rate-limiting enzyme, determined by the apparent deactivation energy *E*_*d*_, leading to a decline in rate beyond the temperature of peak performance (*T*_*pk*_). In effect, each *E* represents the trait-specific “apparent” activation energy reflecting the emergent, aggregate effect of the activation energies of the multiple underlying biochemical reactions contributing to that trait. Each *B*_0_ is a trait-specific normalization constant that represents the trait value at a biologically meaningful reference temperature (*T*_*r*_) within the operational temperature range (Fig. 1) and includes the scaling effects of cell size on diffusion-limited cellular kinetics. Although the Sharpe-Schoolfield model focuses exclusively on enzyme kinetics, any other TPC model can be used to derive the temperature dependence of pairwise interactions using the approach we show below. In section S3, we show that our particular TPC parameterization implicitly includes diffusion limitation as an additional (but secondary) constraint.

### 2.2 Calculating effective species interactions

To derive the pairwise species interactions mediated by the temperature-dependent trait-driven resource utilization dynamics, we obtain the MiCRM’s effective Lotka–Volterra (ELV) system by performing a multivariate Taylor expansion of the consumer dynamics around the quasisteady state of resource abundances, assuming resource dynamics reach equilibrium faster than consumer dynamics (MacArthur 1970, Marsland et al. 2020*a*; section S1):

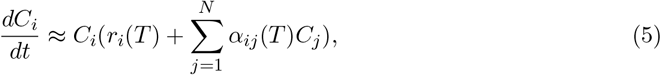

where:

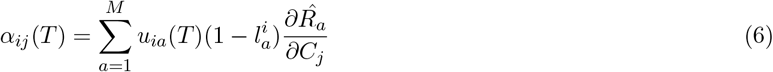

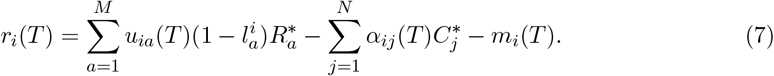

And the effective interaction matrix is:

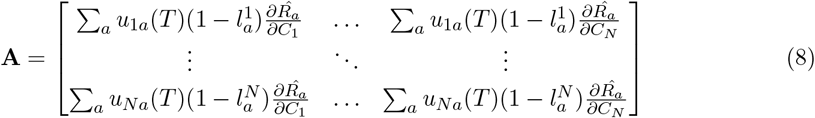

Thus, each effective pairwise species interaction embodies the manipulation of available resources by populations, including the metabolic transformation and leakage by all species within the community (because the stoichiometric matrix of each species is fully connected; Fig. 1), modulated by the equilibrium abundances of both consumers (**C**^∗^) and resources (**R**^∗^). Specifically: (i) The impact of species *j* on *i* is determined by the perturbation on resource equilibrium as the former’s population grows 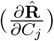, mediated by the preferential, temperature-dependent resource uptake rate of species *i* (*u*_*ia*_(*T* )); (ii) Although the model is formally pairwise (the Taylor expansion is first-order), these coefficients implicitly capture higher-order interactions the aggregate uptake and leakage of all species in the entire community through the inverse because the sensitivity of resources to species 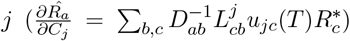 depends on matrix **D** ^−1^, where 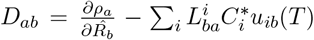 (section S1); (iii) The impact of a species on itself *α*_*ii*_(*T* ) has a similar functional form to *α*_*i*≠*j*_(*T* ), which also encapsulates the net effects of competition and cross-feeding from all species in the community; (iv) The *i*^*th*^ species’ maximal growth rate *r*_*i*_(*T* ) (the theoretical growth rate when species’ abundances are low and resources are effectively unlimited) is its net assimilation of resources accounting for the effects of resource-mediated pairwise interactions minus its (density-independent) respiratory losses (Eqn. 7).

### 2.3 Assembly Simulations

We assembled 999 replicate communities at fixed temperatures between 0–38^◦^C. At each temperature, a random set of 100 consumers and 50 carbon resources was drawn from a global pool of species with varying TPCs for resource uptake and respiration (Fig. 1c), with the assembly dynamics governed by Eqns 1&2.

Each species’ resource uptake rate (*u*_*i*_) was subdivided randomly across all *M* resources (the *u*_*ia*_’s) by sampling resource uptake preferences (proportions) from a Dirichlet distribution (Fig. 1). This allows for the sampling of these proportions from uniform distributions (or beta distributions, permitting different levels of niche overlap depending on parameterization; section S9), and guarantees that the preferential uptake proportions of each species sum to one. Multiplying these proportions by the species-level uptake rate **u**_**i**_ yields the uptake matrix **U**, which describes the preferential uptake rates of each species for each resource.

The leakage-transformation tensor 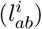, where metabolic interactions determining leakage and the transformation of consumed resources differ for each species, was also sampled from a Dirichlet distribution, such that the sum of the inefficiency of species-level resource utilization on each resource 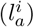 was 0.3 (Fig. 1).

Reflecting the available data on bacterial TPCs, we assumed that the thermal maxima (*T*_*pk*_s) of most species’ traits peaked after 30^◦^C (representing species adapted to mesophilic temperatures; Fig. 2c), with respiration peaking after uptake for any given species (Corkrey et al. 2016, Smith et al. 2019). As such, our framework can be applied to strictly thermophilic or psychrophilic pools of species, provided sufficient trait TPC data are available.

**Figure 2.**
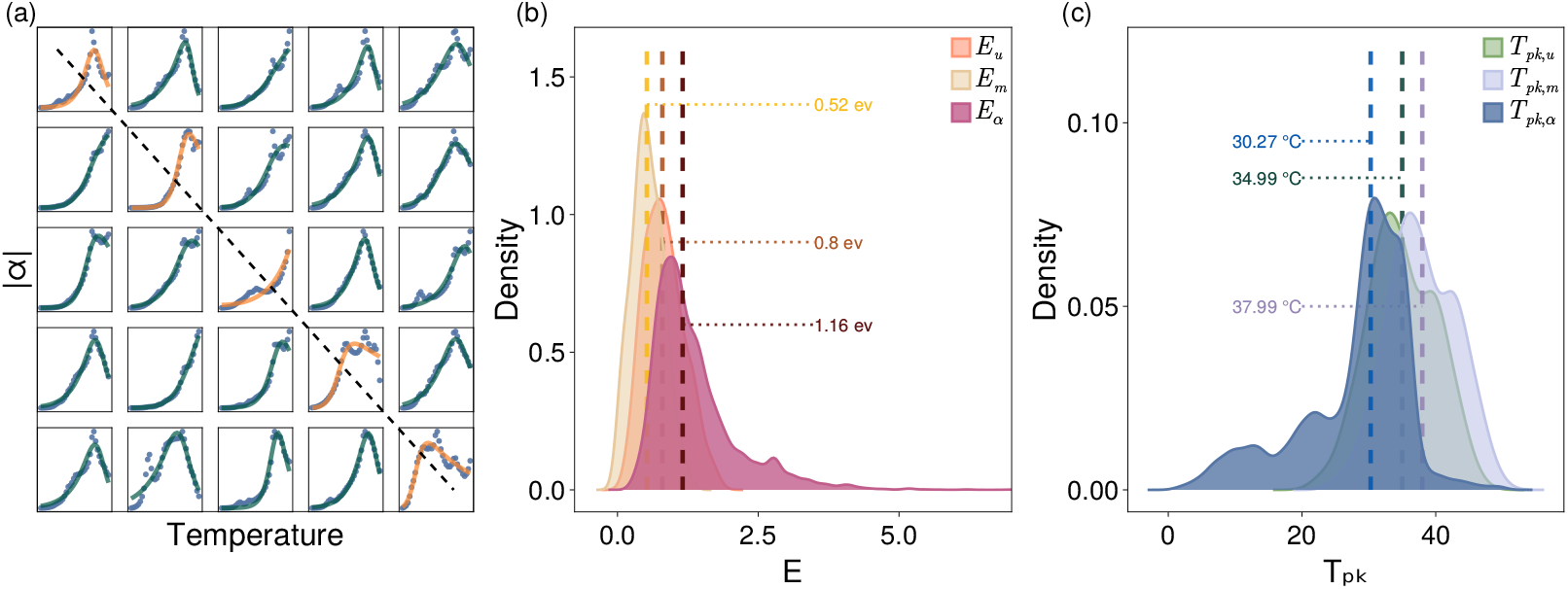
The temperature performance curves (TPCs) of effective interactions in model communities. All calculations were made over 0–38^◦^C. (a) Representative TPCs of intra- and interspecific interaction coefficients (sub-set of the coefficients of one of the 100-species communities simulated in Fig. 3). Orange curves on the diagonal are TPCs of intraspecific interactions (*α*_*ii*_) and off-diagonal green curves the corresponding interspecific interactions (*α*_*i*≠=*j*_). (b) Apparent activation energies of effective interactions (*E*_*α*_; pink). (c) Thermal maxima of interactions (*T*_*pk,α*_; dark blue). The dashed lines mark median values of distributions.

Simulations were implemented in Julia (Bezanson et al. 2017). An example assembly simulation is shown in SI (Fig. S1), and the parameter values used across all simulations are listed in Table S1.

### 2.4 TPC parameterizations

Each species in the immigration pool was assigned randomly generated TPCs for the two traits (*u* and *m*; Eqn. 3 and 4) (Table S2). For this, we sampled each species-specific *B*_0_ and *E* pair (for each of the two traits) from a multivariate normal distribution:

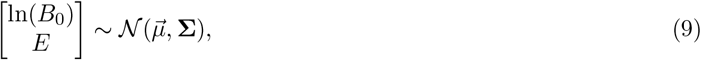

where,

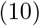

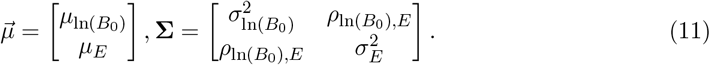

We parameterised the model for simulation using data from 604 TPCs on growth rates and metabolic fluxes, representative of the bacterial tree of life (Smith et al. 2019, 2021). This global dataset synthesizes digitized published experimental curves from diverse taxa spanning the full range of inhabited thermal niches, providing standardized thermodynamic parameters extracted via Sharpe–Schoolfield model fitting.

Based on this dataset, we set 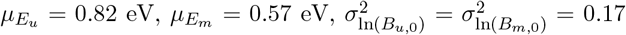 and 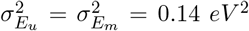. Crucially, the parameterisation 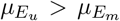 reflects the general empirical pattern published in global microbial datasets, where the thermal sensitivity of growth rates (driven by resource uptake) is typically higher than that of metabolic fluxes (Smith et al. 2019). The correlation coefficient 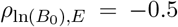, which encodes an intermediate level of thermal generalist-specialist trade-off as observed in empirical data (section S4), was assumed to be identical for the two traits.

Because empirical data on the TPCs of resource uptake rates in bacteria (and microbes in general) are largely absent, we assumed that the distribution of uptake rate thermal sensitivities (*E*_*u*_s) is identical to that of maximal growth rates (*r*_*max*_s). That is, we used the mean and variance of the *E* distribution of species’ *r*_*max*_s as a proxy for 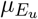 and 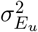. This is based on the fact that anabolic processes (e.g., protein synthesis) underlying bacterial *r*_*max*_ are mainly constrained by resource uptake (Kempes et al. 2012, 2017, Knapp & Huang 2022). Note that *r*_*max*_ here is the exponential growth of species incubated in isolation whereas the *r*_*i*_ derived from the ELV includes the effects of the focal species’ resource-mediated interactions with others (Eqn.7).

Next, 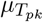 for uptake rates 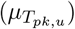 were randomly sampled from a normal distribution. This was done such that the 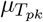 for respiration rate of a given species was always higher than that of uptake (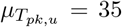 and 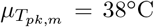). This offset 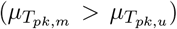 captures a broad physiological phenomenon observed in empirical studies, where catabolic processes (respiration) peak at higher temperatures than anabolic processes (photosynthesis or resource uptake) (Padfield et al. 2016, Smith et al. 2021).

Finally, the normalization constant 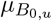 was parameterized by constraining its values to the median value of CUE (species’ carbon use efficiency) at *T*_*r*_ (Smith et al. 2021), *where:*

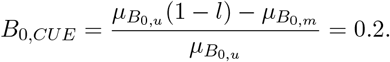

*Because B*_0_ follows a log-normal distribution (Fig.S2 a), 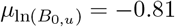 and 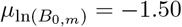. This constraint ensures that species’ respiration rates cannot exceed their resource uptake rates.

The above parameterizations can be applied to other microbial groups by appropriately adjusting values of the TPC parameters (Kontopoulos et al. 2020, Smith et al. 2019).

### 2.5 Quantifying the TPCs of effective interactions

To quantify the temperature dependence of the effective pairwise species interactions, we again fitted the effective coefficients (*α*) to the Sharpe-Schoolfield model:

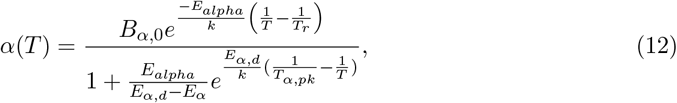

using non-linear least squares with `curve_fit` from the `LsqFit` package in Julia.

### 2.6 Quantifying Richness and Local Stability

Species richness was quantified as the number of surviving species (species with non-zero biomass) at equilibrium. Local stability was quantified by evaluating the community (Jacobian) matrix at the system’s equilibrium and computing the probability that all eigenvalues have negative real parts, indicating that the equilibrium is locally (asymptotically) stable. This probability of stability was calculated as the proportion of locally stable communities across all 999 simulations at each temperature.

## 3 Results

### 3.1 Interactions are more Thermally Sensitive than Underlying Metabolic Traits

Both intraspecific and interspecific interactions have the same TPC form because they emerge from the same resource-mediated mechanisms (Eqn. 8; Fig.2 a). The emergent TPCs of effective pairwise interactions are typically unimodal (Fig. 2a; Fig. S11-S14), peaking earlier (median ≈ 30^◦^C) than the underlying metabolic traits (*T*_*pk,α*_ *< T*_*pk,u*_ *< T*_*pk,m*_; Fig. 2c). This early peak in interaction strength arises from two mechanisms. First, there is a decline in equilibrium abundances at high temperatures: while uptake rates continue to rise with warming, equilibrium abundances of both consumers and resources decrease exponentially at high temperatures (Fig. S4). Since these equilibrium abundances govern the effective interaction coefficients (Eqn. S8), this decline effectively weakens interaction strengths well before uptake rates peak. Second, as temperatures increase, a growing proportion of species’ uptake rate TPCs approach their peak performance (Fig. 2c) and begin to decline.

The effective interactions also typically exhibit higher apparent activation energies (median *E*_*α*_ = 1.16 eV) than those of the underlying metabolic traits (Fig. 2b,c; *E*_*α*_ *> E*_*u*_, *E*_*m*_). This amplification of thermal sensitivity arises from the multiplicative nature of resource-mediated effective interaction coefficients (Eqn. 6; section S2).

Note that respiration TPCs contribute only indirectly to the temperature dependence of species interactions by regulating equilibrium consumer abundances via effective growth rates (Eqn. 7; SI Eqns. SI.14–15), and therefore exert a relatively minor influence on the strength of resource-mediated interactions compared to resource uptake.

Taken together, these results predict that species interactions are intrinsically more sensitive to increasing temperatures than the underlying metabolic traits, both in terms of thermal maxima and activation energies.

### 3.2 Temperature Systematically Shifts the Distributions of Effective Interactions

The distributions of both intraspecific and interspecific interactions are approximately lognormal at all temperatures (Fig. 3). This is because the interaction coefficients emerge from sums of multiplicative terms involving species’ uptake rates, which are themselves log-normally distributed (Eqn. S8). The interactions in these systems are predominantly negative (Fig. 3c) because effective pairwise coefficients are dominated by resource depletion, so shared resource use produces net competitive effects even in the presence of cross-feeding. This is a general property of effective Lotka–Volterra systems derived from consumer–resource dynamics, where pairwise interactions encode the indirect effects of competition for limiting resources (MacArthur 1970, Marsland et al. 2020*a*, Butler & O’Dwyer 2018, Mustri et al. 2025). As temperature increases, resource uptake rates rise exponentially, strengthening this effective competition among species (Eqn. 6; Fig. 3).

**Figure 3.**
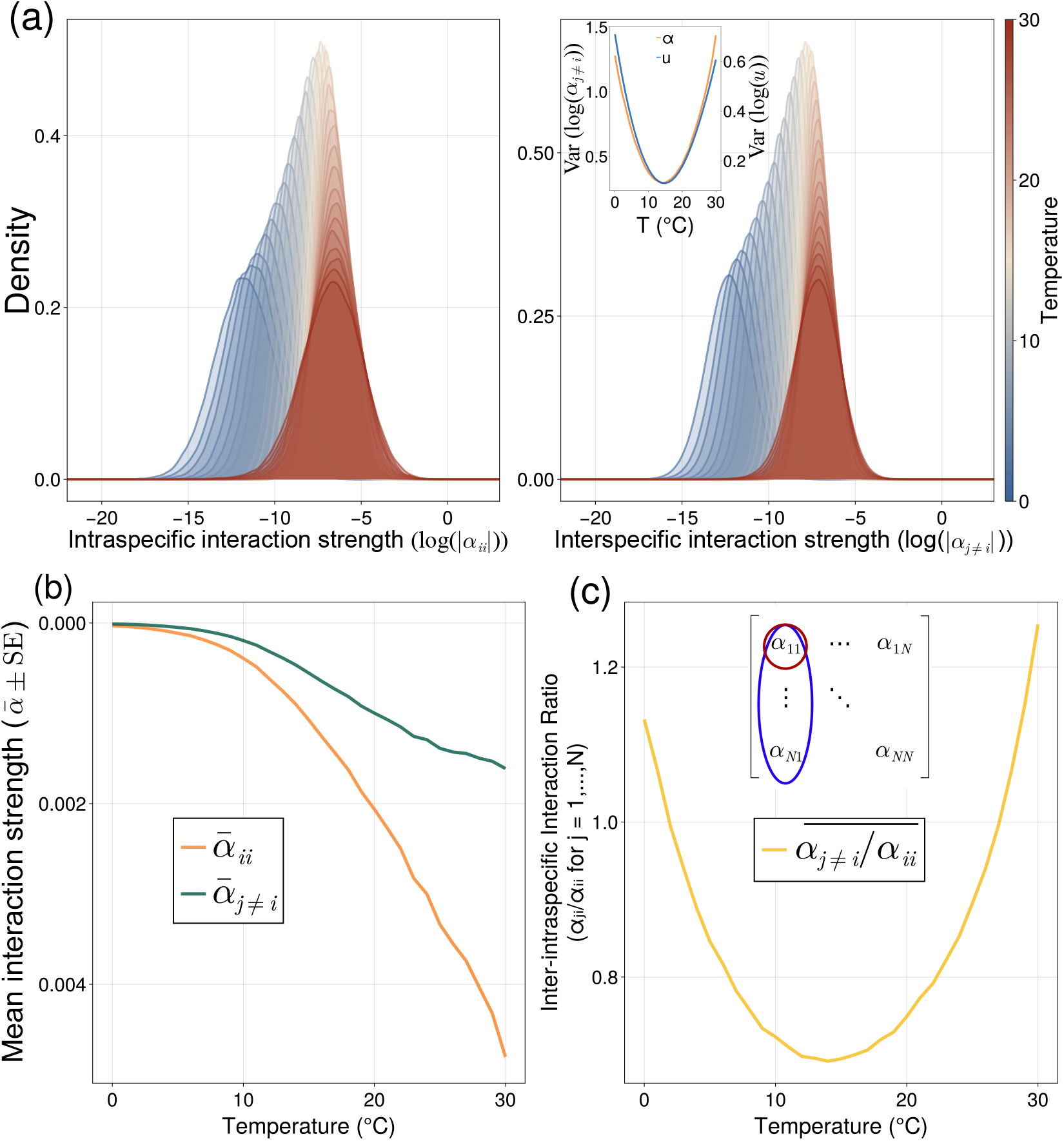
The effect of temperature on effective interaction strengths. **(a)** The distributions of intra- and interspecific interactions are log-normal with variance minimised at intermediate temperature (inset figure), and (**(b)**) mean increasing monotonically with temperature such that intraspecific interactions increase in strength with temperature faster than interspecific interactions. **(c)** The mean of the ratio of each species’ interspecific interactions relative to its intraspecific interaction (self-regulation) is also minimised at intermediate temperature. These results are from 999 model communities at each temperature.

Furthermore, the variance of interaction coefficients is minimized at intermediate temperatures where variation in uptake rates is lowest (Fig. 3a, b). Since effective interactions are constructed from the multiplications of species’ uptake rates (Eqn. 6), reduced variation in uptake rates directly leads to reduced variation in both intra- and interspecific interaction strengths.

The mean absolute values of all interactions increase exponentially with temperature, with intraspecific interactions *α*_*ii*_ increasing more rapidly than interspecific interactions *α*_*i*≠=*j*_ (Fig. 3a). This arises from the log-normality of the interaction coefficients (section S2; each *α*_*ii*_ contains the square of a single species’ uptake rate, whereas *α*_*i*≠=*j*_ contains only the product of two species’ uptake rates).

Altogether, these temperature-dependent patterns in the statistical properties of the effective interaction have important consequences for community dynamics, as we show next.

### 3.3 Consequences for Temperature-dependent Community Dynamics

Species richness follows a unimodal temperature response (Fig. 4a), driven by the fact that effective interactions exhibit minimal variation at intermediate temperatures (Fig. 3b-c) (Clegg&Pawar 2024). Stability is also maximized at intermediate temperatures, where the ratio of inter- to intra-specific interaction strengths is minimized (Fig. 4b; section S2).

**Figure 4.**
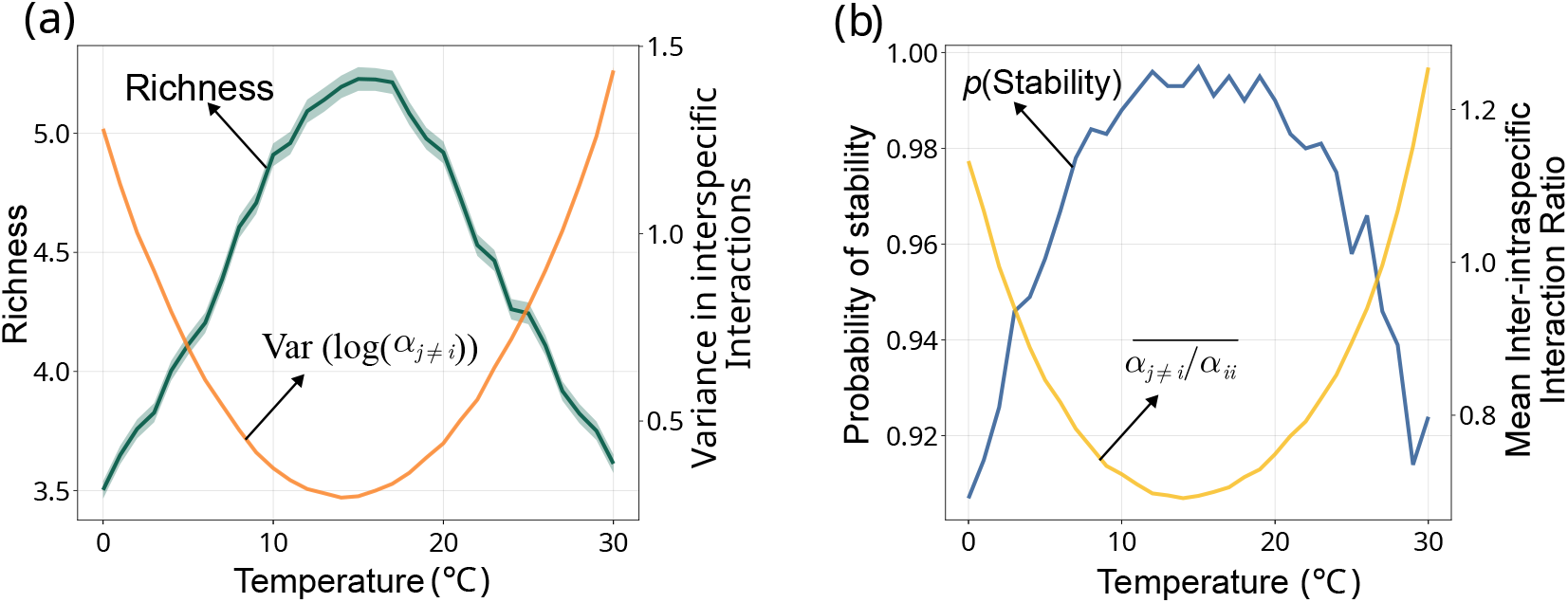
The temperature-dependence of feasibility and stability. (a) Mean species richness (*±*SE) changes unimodally with temperature, peaking where variation in interspecific interaction strengths is minimized (Fig. 3a). (b) Probability of stability is maximised at intermediate temperatures, inter– to intraspecific interaction strength ratio is minimized (Fig. 3c). Data are from the same simulations as in Fig. 3.

## 4 Discussion

By explicitly capturing the combined temperature-dependence of resource competition, facilitation, and cross-feeding in pairwise consumer species interaction coefficients, our results provide a basis for mechanistic understanding and for deriving temperature-dependent microbial species interactions, and how these interactions, in turn, shape community structure, diversity, and stability.

That effective species interactions are typically more thermally sensitive than the underlying metabolic traits, both in terms of apparent activation energy (*E*) and temperature of peak performance (*T*_*pk*_) (Fig. 2) has two corresponding implications. Firstly, the higher *E* of species interactions (relative to metabolic traits) means that overall competitiveness in the community increases faster than would be expected from the underlying metabolic traits alone (Fig. 2a-b). Therefore, community diversity should increase at a lower rate towards intermediate temperatures than expected from the TPCs of metabolic traits alone (Clegg & Pawar 2024). Secondly, the lower temperatures of peak interaction strengths relative to those of underlying metabolic traits means that beyond a critical temperature (*approx*30^◦^C in the case of mesophilic bacterial communities; Fig. 2c)), changes in community dynamics with further warming would be modulated by declines in interaction strengths. In general, this suggests that the sensitivity of community-level dynamics to gradual temperature change (e.g., global warming) or gradients (e.g., latitudinal or altitudinal) may be greater than predicted from the TPCs of physiological traits alone.

Beyond the thermal sensitivity of species interactions, our results show that temperature fundamentally reshapes their statistical distributions (Fig. 3). Firstly, variation in effective pairwise interactions reaches a minimum at intermediate temperatures (Fig. 3). This is where the feasibility of community equilibria is maximized (Clegg & Pawar 2024), resulting in a unimodal response of species richness to temperature (Fig. 4a). Secondly, the ratio of interspecific to intraspecific interaction strength, *α*_*j i*_*/α*_*ii*_, is also minimized at intermediate temperatures (Fig. 4b; section S2). This is where the diagonal dominance (effective self-regulation) of the community (Jacobian) matrix is maximized (Haydon 1994, Barabás et al. 2017), resulting in the maximization of the probability of local, asymptotic stability (Fig. 4b). The effective selfregulation ratio *α*_*j*≠ *i*_*/α*_*ii*_ is a measure of relative competitiveness between species pairs (Chesson 2000) and quantifies the likelihood of competitive exclusion: when intraspecific interactions dominate over interspecific ones, exclusion is less likely, and coexistence is favored. Therefore, species richness peaks at an intermediate temperature (Fig. 4), where antagonism among species is weakest. The unimodality of this ratio also implies that, at relatively low and high temperatures, species effectively spend more metabolic energy on resource exploitation than on effective self-regulation.

In section S7 and S10, we show that our findings about the characteristics of the temperature dependence of effective interactions and their consequences for community dynamics hold across diverse ecological contexts, from well-mixed microbial communities with cross-feeding to classical competitive consumer–resource systems without leakage under multiple environmentally realistic resource supply regimes. These results are also qualitatively robust to the inclusion or exclusion of cross-feeding, as removing metabolic leakage does not qualitatively alter the temperature dependence of effective interactions (section S2 and section S7). Indeed, our framework generalizes further, to pairwise interactions across trophic levels as long as a timescale separation between resource and consumer dynamics exists. This separation is expected to hold not only for bacteria and substrates, but also for higher trophic levels (e.g., protists consuming bacteria) because mass-specific biological rates scale negatively with body size, and the same underlying traits at different trophic levels—consumption and respiration—follow the same fundamental thermodynamic constraints. Generalizing our approach to autotrophs such as phytoplankton would require fundamental changes to the consumer-resource model to include light and nutrient limitation, and is a promising avenue for future work.

Empirical validation of these predictions requires time-series data of species and metabolite abundances across a thermal gradient, supported by direct measurements of uptake and respiration rates. Fitting dynamical models to such data would allow for the rigorous inference of interaction thermal performance curves. Although these comprehensive datasets are currently unavailable, our framework provides a clear path forward to bridge this gap by explicitly linking these community-level dynamics to measurable physiological traits.

Lacking sufficient data on the temperature dependence of uptake rates, we used the activation energies of species’ *r*_*max*_s to parameterize resource uptake rate TPCs. Rapid advances in technologies such as microfluidics and isotope-based methods are likely to make measurements of microbial resource uptake increasingly feasible at scale, thereby improving the accuracy of predictions in future studies that apply this framework. Indeed, as multiomics technologies continue to advance, our framework offers a promising route to predicting interaction structure directly from species’ intrinsic metabolic properties. For more realistic predictions of community dynamics, Genome Scale Metabolic Models (GEMs) (Wang 2017) could be used to map the underlying biochemical networks and enzyme kinetics of cellular-level metabolism to provide parameterizations for the leakage tensor and resource uptake matrices, and provide a mechanistic understanding of competitive and facilitative interactions through metabolic pathways and flux exchanges (Zelezniak et al. 2015, Machado et al. 2021).

Finally, we acknowledge that additional factors, including predation (Pérez et al. 2016), quorum sensing (Miller & Bassler 2001, Whitehead et al. 2001), horizontal gene transfer (Arnold et al. 2022), cannibalism (González-Pastor et al. 2003), chemical inhibition, and competition for limiting nutrients such as nitrogen and phosphorus (Koide et al. 2005) may modify the temperature-dependence of pairwise microbial interactions and their community-level consequences. Identifying which of these factors (or their combinations) play a key role in specific environmental or biological contexts remains a promising and necessary avenue for future work.

## 5 Code Availability

Code for simulation and analysis in this study is available at https://github.com/DaniDuan/temp_interactions.git.

## References

Arnold, B. J., Huang, I.-T. & Hanage, W. P. (2022), ‘Horizontal gene transfer and adaptive evolution in bacteria’, Nature Reviews Microbiology 20(4), 206–218.

Arnoldi, J.-F., Jackson, A. L., Peralta-Maraver, I. & Payne, N. L. (2025), ‘A universal thermal performance curve arises in biology and ecology’, Proceedings of the National Academy of Sciences 122(43), e2513099122. URL: https://www.pnas.org/doi/abs/10.1073/pnas.2513099122

Bar-On, Y. M., Phillips, R. & Milo, R. (2018), ‘The biomass distribution on earth’, Proceedings of the National Academy of Sciences 115(25), 6506–6511.

Barabás, G., Michalska-Smith, M. J. & Allesina, S. (2017), ‘Self-regulation and the stability of large ecological networks’, Nature ecology & evolution 1(12), 1870–1875.

Barberán, A., Bates, S. T., Casamayor, E. O. & Fierer, N. (2012), ‘Using network analysis to explore co-occurrence patterns in soil microbial communities’, The ISME journal 6(2), 343–351.

Bestion, E., García-Carreras, B., Schaum, C.-E., Pawar, S. & Yvon-Durocher, G. (2018), ‘Metabolic traits predict the effects of warming on phytoplankton competition’, Ecology Letters 21(5), 655–664.

Bezanson, J., Edelman, A., Karpinski, S. & Shah, V. B. (2017), ‘Julia: A fresh approach to numerical computing’, SIAM Review 59(1), 65–98. URL: https://epubs.siam.org/doi/10.1137/141000671

Blanchet, F. G., Cazelles, K. & Gravel, D. (2020), ‘Co-occurrence is not evidence of ecological interactions’, Ecology Letters 23(7), 1050–1063.

Butler, S. & O’Dwyer, J. P. (2018), ‘Stability criteria for complex microbial communities’, Nature communications 9(1), 2970.

Cazelles, K., Araújo, M. B., Mouquet, N. & Gravel, D. (2016), ‘A theory for species cooccurrence in interaction networks’, Theoretical Ecology 9, 39–48.

Chesson, P. (2000), ‘Mechanisms of maintenance of species diversity’, Annual review of Ecology and Systematics 31(1), 343–366.

Clegg, T. & Pawar, S. (2024), ‘Variation in thermal physiology can drive the temperature-dependence of microbial community richness’, eLife 13, e84662. URL: 10.7554/eLife.84662

Corkrey, R., McMeekin, T. A., Bowman, J. P., Ratkowsky, D. A., Olley, J. & Ross, T. (2016), ‘The biokinetic spectrum for temperature’, PLoS One 11(4), e0153343.

Corkrey, R., Olley, J., Ratkowsky, D., McMeekin, T. & Ross, T. (2012), ‘Universality of ther-modynamic constants governing biological growth rates’, PLoS One 7(2), e32003.

Cui, W., Marsland, R. & Mehta, P. (2024), ‘Les houches lectures on community ecology: From niche theory to statistical mechanics’, ArXiv pp. arXiv–2403.

De Baar, H. J. W. v. (1994), ‘von liebig’s law of the minimum and plankton ecology (1899– 1991)’, Progress in oceanography 33(4), 347–386.

Dell, A. I., Pawar, S. & Savage, V. M. (2011), ‘Systematic variation in the temperature dependence of physiological and ecological traits’, Proceedings of the National Academy of Sciences 108(26), 10591–10596.

Dell, A. I., Pawar, S. & Savage, V. M. (2014), ‘Temperature dependence of trophic interactions are driven by asymmetry of species responses and foraging strategy’, Journal of Animal Ecology 83(1), 70–84.

DeLong, J. P., Okie, J. G., Moses, M. E., Sibly, R. M. & Brown, J. H. (2010), ‘Shifts in metabolic scaling, production, and efficiency across major evolutionary transitions of life’, Proceedings of the National Academy of Sciences 107(29), 12941–12945.

García, F. C., Clegg, T., O’Neill, D. B., Warfield, R., Pawar, S. & Yvon-Durocher, G. (2023), ‘The temperature dependence of microbial community respiration is amplified by changes in species interactions’, Nature microbiology 8(2), 272–283.

Gibbs, T., Zhang, Y., Miller, Z. R. & O’Dwyer, J. P. (2022), ‘Stability criteria for the consumption and exchange of essential resources’, PLoS computational biology 18(9), e1010521.

Gilbert, B., Tunney, T. D., McCann, K. S., DeLong, J. P., Vasseur, D. A., Savage, V., Shurin, J. B., Dell, A. I., Barton, B. T., Harley, C. D. et al. (2014), ‘A bioenergetic framework for the temperature dependence of trophic interactions’, Ecology letters 17(8), 902–914.

Goldford, J. E., Lu, N., Bajić, D., Estrela, S., Tikhonov, M., Sanchez-Gorostiaga, A., Segre, D., Mehta, P. & Sanchez, A. (2018), ‘Emergent simplicity in microbial community assembly’, Science 361(6401), 469–474.

González-Pastor, J. E., Hobbs, E. C. & Losick, R. (2003), ‘Cannibalism by sporulating bacteria’, Science 301(5632), 510–513.

Handley, K. M. (2019), ‘Determining microbial roles in ecosystem function: redefining microbial food webs and transcending kingdom barriers’, Msystems 4(3), e00153–19.

Haydon, D. (1994), ‘Pivotal assumptions determining the relationship between stability and complexity: an analytical synthesis of the stability-complexity debate’, The American Naturalist 144(1), 14–29.

Kaur, T. & Dutta, P. S. (2020), ‘Persistence and stability of interacting species in response to climate warming: the role of trophic structure’, Theoretical Ecology 13(3), 333–348.

Kempes, C. P., Dutkiewicz, S. & Follows, M. J. (2012), ‘Growth, metabolic partitioning, and the size of microorganisms’, Proceedings of the National Academy of Sciences 109(2), 495–500.

Kempes, C. P., van Bodegom, P. M., Wolpert, D., Libby, E., Amend, J. & Hoehler, T. (2017), ‘Drivers of bacterial maintenance and minimal energy requirements’, Frontiers in microbiology 8, 31.

Klausmeier, C. A., Litchman, E. & Levin, S. A. (2004), ‘Phytoplankton growth and stoichiometry under multiple nutrient limitation’, Limnology and oceanography 49(4part2), 1463–1470.

Knapp, B. D. & Huang, K. C. (2022), ‘The Effects of Temperature on Cellular Physiology’, Annu. Rev. Biophys. 51(1), 499–526.

Koide, R. T., Xu, B., Sharda, J., Lekberg, Y. & Ostiguy, N. (2005), ‘Evidence of species interactions within an ectomycorrhizal fungal community’, New Phytologist pp. 305–316.

Kontopoulos, D., Smith, T. P., Barraclough, T. G. & Pawar, S. (2020), ‘Adaptive evolution shapes the present-day distribution of the thermal sensitivity of population growth rate’, PLoS biology 18(10), e3000894.

Lax, S., Abreu, C. I. & Gore, J. (2020), ‘Higher temperatures generically favour slower-growing bacterial species in multispecies communities’, Nat. Ecol. Evol. 4(4), 560–567.

Lechón-Alonso, P., Clegg, T., Cook, J., Smith, T. P. & Pawar, S. (2021), ‘The role of competition versus cooperation in microbial community coalescence’, PLOS Computational Biology 17(11), e1009584.

MacArthur, R. (1970), ‘Species packing and competitive equilibrium for many species’, Theoretical population biology 1(1), 1–11.

MacArthur, R. M. (1969), ‘Species packing, and what competition minimizes’, Proceedings of the National Academy of Sciences 64(4), 1369–1371.

Machado, D., Maistrenko, O. M., Andrejev, S., Kim, Y., Bork, P., Patil, K. R. & Patil, K. R. (2021), ‘Polarization of microbial communities between competitive and cooperative metabolism’, Nature ecology & evolution 5(2), 195–203.

Malik, A. A., Martiny, J. B., Brodie, E. L., Martiny, A. C., Treseder, K. K. & Allison, S. D. (2020), ‘Defining trait-based microbial strategies with consequences for soil carbon cycling under climate change’, The ISME journal 14(1), 1–9.

Marsland, R., Cui, W., Goldford, J., Sanchez, A., Korolev, K. & Mehta, P. (2019), ‘Available energy fluxes drive a transition in the diversity, stability, and functional structure of microbial communities’, PLoS computational biology 15(2), e1006793.

Marsland, R., Cui, W. & Mehta, P. (2020a), ‘A minimal model for microbial biodiversity can reproduce experimentally observed ecological patterns’, Scientific reports 10(1), 1–17.

Marsland, R., Cui, W. & Mehta, P. (2020b), ‘The minimum environmental perturbation principle: A new perspective on niche theory’, The American Naturalist 196(3), 291–305.

Miller, M. B. & Bassler, B. L. (2001), ‘Quorum sensing in bacteria’, Annual Reviews in Microbiology 55(1), 165–199.

Mustri, M. P., Duan, Q. & Pawar, S. (2025), ‘Accuracy of the lotka-volterra model fails in strongly coupled microbial consumer-resource systems’, PLOS Computational Biology 21(12), 1–16. URL: 10.1371/journal.pcbi.1013719

Padfield, D., Yvon-Durocher, G., Buckling, A., Jennings, S. & Yvon-Durocher, G. (2016), ‘Rapid evolution of metabolic traits explains thermal adaptation in phytoplankton’, Ecology letters 19(2), 133–142.

Pérez, J., Moraleda-Muñoz, A., Marcos-Torres, F. J. & Muñoz-Dorado, J. (2016), ‘Bacterial predation: 75 years and counting!’, Environmental Microbiology 18(3), 766–779.

Philippot, L., Spor, A., Hénault, C., Bru, D., Bizouard, F., Jones, C. M., Sarr, A. & Maron, P.-A. (2013), ‘Loss in microbial diversity affects nitrogen cycling in soil’, The ISME journal 7(8), 1609–1619.

Pold, G., Domeignoz-Horta, L. A., Morrison, E. W., Frey, S. D., Sistla, S. A. & DeAngelis, K. M. (2020), ‘Carbon use efficiency and its temperature sensitivity covary in soil bacteria’, MBio 11(1), 10–1128.

Rivett, D. W. & Bell, T. (2018), ‘Abundance determines the functional role of bacterial phylotypes in complex communities’, Nature microbiology 3(7), 767–772.

Schoolfield, R. M., Sharpe, P. & Magnuson, C. E. (1981), ‘Non-linear regression of biological temperature-dependent rate models based on absolute reaction-rate theory’, Journal of theoretical biology 88(4), 719–731.

Smith, T. P., Clegg, T., Bell, T. & Pawar, S. (2021), ‘Systematic variation in the temperature dependence of bacterial carbon use efficiency’, Ecology Letters 24(10), 2123–2133.

Smith, T. P., Thomas, T. J., García-Carreras, B., Sal, S., Yvon-Durocher, G., Bell, T. & Pawar, S. (2019), ‘Community-level respiration of prokaryotic microbes may rise with global warming’, Nature communications 10(1), 5124.

Stark, K. A., Clegg, T., Bernhardt, J. R., Grainger, T. N., Kempes, C. P., Savage, V., O’Connor, M. I. & Pawar, S. (2024), ‘Towards a more dynamic metabolic theory of ecology to predict climate change effects on biological systems’.

The Human Microbiome Project Consortium, Huttenhower, C., Gevers, D., Knight, R., Abubucker, S., Badger, J. H., Chinwalla, A. T., Creasy, H. H., Earl, A. M., Fitzgerald, M. G., Fulton, R. S., Giglio, M. G., Hallsworth-Pepin, K., Lobos, E. A., Madupu, R., Magrini, V., Martin, J. C., Mitreva, M., Muzny, D. M., Sodergren, E. J., Versalovic, J., Wollam, A. M., Worley, K. C., Wortman, J. R., Young, S. K., Zeng, Q., Aagaard, K. M., Abolude, O. O., Allen-Vercoe, E., Alm, E. J., Alvarado, L., Andersen, G. L., Anderson, S., Appelbaum, E., Arachchi, H. M., Armitage, G., Arze, C. A., Ayvaz, T., Baker, C. C., Begg, L., Belachew, T., Bhonagiri, V., Bihan, M., Blaser, M. J., Bloom, T., Bonazzi, V., Paul Brooks, J., Buck, G. A., Buhay, C. J., Busam, D. A., Campbell, J. L., Canon, S. R., Cantarel, B. L., Chain, P. S. G., Chen, I. M. A., Chen, L., Chhibba, S., Chu, K., Ciulla, D. M., Clemente, J. C., Clifton, S. W., Conlan, S., Crabtree, J., Cutting, M. A., Davidovics, N. J., Davis, C. C., Desantis, T. Z., Deal, C., Delehaunty, K. D., Dewhirst, F. E., Deych, E., Ding, Y., Dooling, D. J., Dugan, S. P., Michael Dunne, W., Scott Durkin, A., Edgar, R. C., Erlich, R. L., Farmer, C. N., Farrell, R. M., Faust, K., Feldgarden, M., Felix, V. M., Fisher, S., Fodor, A. A., Forney, L. J., Foster, L., di Francesco, V., Friedman, J., Friedrich, D. C., Fronick, C. C., Fulton, L. L., Gao, H., Garcia, N., Giannoukos, G., Giblin, C., Giovanni, M. Y., Goldberg, J. M., Goll, J., Gonzalez, A., Griggs, A., Gujja, S., Kinder Haake, S., Haas, B. J., Hamilton, H. A., Harris, E. L., Hepburn, T. A., Herter, B., Hoffmann, D. E., Holder, M. E., Howarth, C., Huang, K. H., Huse, S. M., Izard, J., Jansson, J. K., Jiang, H., Jordan, C., Joshi, V., Katancik, J. A., Keitel, W. A., Kelley, S. T., Kells, C., King, N. B., Knights, D., Kong, H. H., Koren, O., Koren, S., Kota, K. C., Kovar, C. L., Kyrpides, N. C., La Rosa, P. S., Lee, S. L., Lemon, K. P., Lennon, N., Lewis, C. M., Lewis, L., Ley, R. E., Li, K., Liolios, K., Liu, B., Liu, Y., Lo, C.-C., Lozupone, C. A., Dwayne Lunsford, R., Madden, T., Mahurkar, A. A., Mannon, P. J., Mardis, E. R., Markowitz, V. M., Mavromatis, K., McCorrison, J. M., McDonald, D., McEwen, J., McGuire, A. L., McInnes, P., Mehta, T., Mihindukulasuriya, K. A., Miller, J. R., Minx, P. J., Newsham, I., Nusbaum, C., O’Laughlin, M., Orvis, J., Pagani, I., Palaniappan, K., Patel, S. M., Pearson, M., Peterson, J., Podar, M., Pohl, C., Pollard, K. S., Pop, M., Priest, M. E., Proctor, L. M., Qin, X., Raes, J., Ravel, J., Reid, J. G., Rho, M., Rhodes, R., Riehle, K. P., Rivera, M. C., Rodriguez-Mueller, B., Rogers, Y.-H., Ross, M. C., Russ, C., Sanka, R. K., Sankar, P., Fah Sathirapongsasuti, J., Schloss, J. A., Schloss, P. D., Schmidt, T. M., Scholz, M., Schriml, L., Schubert, A. M., Segata, N., Segre, J. A., Shannon, W. D., Sharp, R. R., Sharpton, T. J., Shenoy, N., Sheth, N. U., Simone, G. A., Singh, I., Smillie, C. S., Sobel, J. D., Sommer, D. D., Spicer, P., Sutton, G. G., Sykes, S. M., Tabbaa, D. G., Thiagarajan, M., Tomlinson, C. M., Torralba, M., Treangen, T. J., Truty, R. M., Vishnivetskaya, T. A., Walker, J., Wang, L., Wang, Z., Ward, D. V., Warren, W., Watson, M. A., Wellington, C., Wetterstrand, K. A., White, J. R., Wilczek-Boney, K., Wu, Y., Wylie, K. M., Wylie, T., Yandava, C., Ye, L., Ye, Y., Yooseph, S., Youmans, B. P., Zhang, L., Zhou, Y., Zhu, Y., Zoloth, L., Zucker, J. D., Birren, B. W., Gibbs, R. A., Highlander, S. K., Methé, B. A., Nelson, K. E., Petrosino, J. F., Weinstock, G. M., Wilson, R. K. & White, O. (2012), ‘Structure, function and diversity of the healthy human microbiome’, Nature 486(7402), 207–214.

Thompson, L. R., Sanders, J. G., McDonald, D., Amir, A., Ladau, J., Locey, K. J., Prill, R. J., Tripathi, A., Gibbons, S. M., Ackermann, G. et al. (2017), ‘A communal catalogue reveals earth’s multiscale microbial diversity’, Nature 551(7681), 457–463.

Vasseur, D. a. & McCann, K. S. (2005), ‘A mechanistic approach for modeling temperature-dependent consumer-resource dynamics.’, Am. Nat. 166(2), 184–198.

Wang, D. (2017), ‘Systems biology: Constraint-based reconstruction and analysis’.

Whitehead, N. A., Barnard, A. M., Slater, H., Simpson, N. J. & Salmond, G. P. (2001), ‘Quorum-sensing in gram-negative bacteria’, FEMS microbiology reviews 25(4), 365–404.

Zelezniak, A., Andrejev, S., Ponomarova, O., Mende, D. R., Bork, P. & Patil, K. R. (2015), ‘Metabolic dependencies drive species co-occurrence in diverse microbial communities’, Proceedings of the National Academy of Sciences 112(20), 6449–6454.

