## Supplementary Information for "General Patterns in the Temperature-Dependence of Heterotrophic Microbial Interactions"

#### S1 Derivation of the effective Lotka-Volterra model

We start with the MiCRM (Eqn. 1 and 2 in Methods) (Table S1):

$$\frac{dC_i}{dt} = C_i \left( \sum_{a=1}^{M} u_{ia}(1 - \sum_{b=1}^{M} l^i_{ab}) R_a - m_i \right), \tag{S1}$$

$$\frac{dR_a}{dt} = \rho_a(\mathbf{R}) - \sum_{i=1}^{N} \Big( C_i u_{ia} R_a - \sum_{b=1}^{M} C_i u_{ib} R_b l^i_{ba} \Big). \tag{S2}$$

Table S1: **Parameters of the MiCRM.**

| Symbols | Definition | Values | Units |
|---|---|---|---|
| $C_i^0$ | Initial biomass abundance of consumer $i$ | 0.1 | mass |
| $R_a^0$ | Initial resource abundance $a$ | 1.0 | mass |
| $u_{ia}$ | Uptake rate of consumer $i$ on resource $a$ | Temperature-dependent | 1/mass*time |
| $m_i$ | Respiration rate of consumer $i$ | Temperature-dependent | 1/time |
| $l^i_{ab}$ | Leakage/transformation fraction of resource $a$ to $b$ by species $i$ | $l^i_a = 0.3$ | fraction |
| $\rho_a$ | External resource supply of resource $a$ | 1.0 | mass/time |

The effective LV (ELV) system of this MiCRM is an approximation of the consumer dynamics (Eqn. S1) assuming quasi-steady-state of the resource abundances (Eqn. S2) (MacArthur 1970, Marsland et al. 2020). That is, assuming that resource abundances equilibrate much faster than those of the consumers.

First, we express the equilibrium resource abundance (Eqn. S2) as an implicit function of the consumer biomass vector $\mathbf{C}$, denoted as $\hat{R}_a(\mathbf{C})$. We then approximate the biomass dynamics (Eqn. S1) around the equilibrium state $\mathbf{C} = \mathbf{C}^*$ using a first-order multivariate

Taylor expansion:

$$\begin{aligned}
\frac{dC_i}{dt} &= C_i\left(\sum_{a=1}^{M} u_{ia}(1-l_a^i)\hat{R}_a(\mathbf{C}) - m_i\right)\\
&\approx C_i\left[\sum_{a=1}^{M} u_{ia}(1-l_a^i)\left(\hat{R}_a + \sum_{j=1}^{N}\left.\frac{\partial \hat{R}_a}{\partial C_j}\right|_{\mathbf{C}=\mathbf{C}^*}(C_j - C_j^*)\right) - m_i\right]\\
&= C_i(r_i + \sum_{j=1}^{N}\alpha_{ij}C_j)
\end{aligned}$$

The implicit function $\hat{R}_a(\mathbf{C})$ is defined by the quasi-steady-state condition $dR_a/dt = 0$. Then, $\hat{\mathbf{R}}$ is the vector of the $\hat{R}_a(\mathbf{C})$s over all resource types $(a = 1, ..., M)$, henceforth denoted as $\hat{\mathbf{R}}$ for simplicity. Using Eqn. S2, this condition is written as:

$$\rho_a(\hat{\mathbf{R}}) - \sum_i\Big(C_i u_{ia}(T)\hat{R}_a - \sum_b C_i u_{ib}(T)\hat{R}_b l_{ba}^i\Big) = 0, \tag{S3}$$

To derive the interaction coefficients, we need to calculate the sensitivity of equilibrium resource abundances $(\mathbf{R} = \mathbf{R}^*)$ to changes in consumer biomass, denoted by the partial derivatives $\left.\frac{\partial \hat{\mathbf{R}}}{\partial C_j}\right|_{\mathbf{C}=\mathbf{C}^*}$. We calculate this by differentiating the steady-state condition with respect to the biomass of a focal species $j$ and applying the chain rule. This yields:

$$\sum_b \frac{\partial \rho_a}{\partial \hat{R}_b}\frac{\partial \hat{R}_b}{\partial C_j} - \left(u_{ja}R_a^* + \sum_i C_i^* u_{ia}\frac{\partial \hat{R}_a}{\partial C_j}\right) + \left(\sum_b u_{jb}R_b^* l_{ba}^j + \sum_i\sum_b C_i^* u_{ib} l_{ba}^i \frac{\partial \hat{R}_b}{\partial C_j}\right) = 0. \tag{S4}$$

Rearranging the equation:

$$\sum_b \left[\frac{\partial \rho_a}{\partial \hat{R}_b} - \sum_i(\delta_{ba} - l_{ba}^i)C_i^* u_{ib}\right]\frac{\partial \hat{R}_b}{\partial C_j} = \sum_b(\delta_{ba} - l_{ba}^j)u_{jb}R_b^*. \tag{S5}$$

For simpler notation, we assign $L_{ba}^i = \delta_{ba} - l_{ba}^i$, where $\delta_{ba}$ represents the Kronecker delta that $\delta_{ab} = \begin{cases} 0, & \text{if } a \neq b \\ 1, & \text{if } a = b \end{cases}$. We also define the matrix elements $D_{ab} = \frac{\partial \rho_a}{\partial \hat{R}_b} - \sum_i L_{ba}^i C_i^* u_{ib}$. This simplifies the system to:

$$\sum_b D_{ab}\frac{\partial \hat{R}_b}{\partial C_j} = \sum_b L_{ba}^j u_{jb} R_b^*.$$

Solving this linear system for the vector of partial derivatives $\frac{\partial \hat{\mathbf{R}}}{\partial C_j}\Big|_{\mathbf{C}=\mathbf{C}^*}$:

$$
\begin{aligned}
\frac{\partial \hat{\mathbf{R}}}{\partial C_j} &= \mathbf{D}^{-1} \cdot \mathbf{L}^j \cdot \mathbf{u}\mathbf{R}^* \qquad \text{(S6)}\\
&= \begin{bmatrix} D^{-1}_{11} & \dots & D^{-1}_{1M} \\ \vdots & \ddots & \\ D^{-1}_{M1} & \dots & D^{-1}_{MM} \end{bmatrix} \begin{bmatrix} L^j_{11} & \dots & L^j_{M1} \\ \vdots & \ddots & \\ L^j_{1M} & \dots & L^j_{MM} \end{bmatrix} \begin{bmatrix} u_{j1}R^*_1 \\ \vdots \\ u_{jM}R^*_M \end{bmatrix}\\
&= \begin{bmatrix} D^{-1}_{11} & \dots & D^{-1}_{1M} \\ \vdots & \ddots & \\ D^{-1}_{M1} & \dots & D^{-1}_{MM} \end{bmatrix} \begin{bmatrix} \sum_c L^j_{c1} u_{jc} R^*_c \\ \vdots \\ \sum_c L^j_{cM} u_{jc} R^*_c \end{bmatrix}\\
&= \begin{bmatrix} \sum_{b,c} D^{-1}_{1b} L^j_{cb} u_{jc} R^*_c \\ \vdots \\ \sum_{b,c} D^{-1}_{Mb} L^j_{cb} u_{jc} R^*_c \end{bmatrix}.
\end{aligned}
$$

The effect of the $j^{th}$ consumer on the $a^{th}$ resource at equilibrium is:

$$\frac{\partial \hat{R}_a}{\partial C_j} = \sum_{b,c} D^{-1}_{ab} L^j_{cb} u_{jc} R^*_c.$$

Thus, the complete equation for the interaction coefficient is:

$$
\begin{aligned}
\alpha_{ij} &= \sum_a u_{ia}(1 - l^i_a)\frac{\partial \hat{R}_a}{\partial C_j} \qquad \text{(S7)}\\
&= \sum_{a,b,c} u_{ia}(1 - l^i_a) D^{-1}_{ab} L^j_{cb} u_{jc} R^*_c. \qquad \text{(S8)}
\end{aligned}
$$

And the effective intrinsic growth rate is:

$$r_i = \sum_a u_{ia}(1 - l^i_a)R^*_a - \sum_j \alpha_{ij} C^*_j - m_i. \qquad \text{(S9)}$$

Note that the pairwise species interaction network ($\mathbf{A}$) that emerges from the MiCRM will always be fully connected because all populations are able to access all resources in a well-mixed environment. Also, during parameterization for simulations, we set the leakage fraction to be 0.3 for every species. Sec. S10 shows that changing leakage levels (Fig. S10), or the uptake structure (Sec. S9, Fig. S9; scenarios of high niche differentiation), does not qualitatively affect our results.

Figure S1 shows example simulations comparing the MiCRM and ELV dynamics at two temperatures. Since the ELV is an approximation of the MiCRM at resource equilibrium, it does not capture the initial transient consumer biomass dynamics in the latter, but converges on the "correct" equilbrium abundances (Fig. S1).

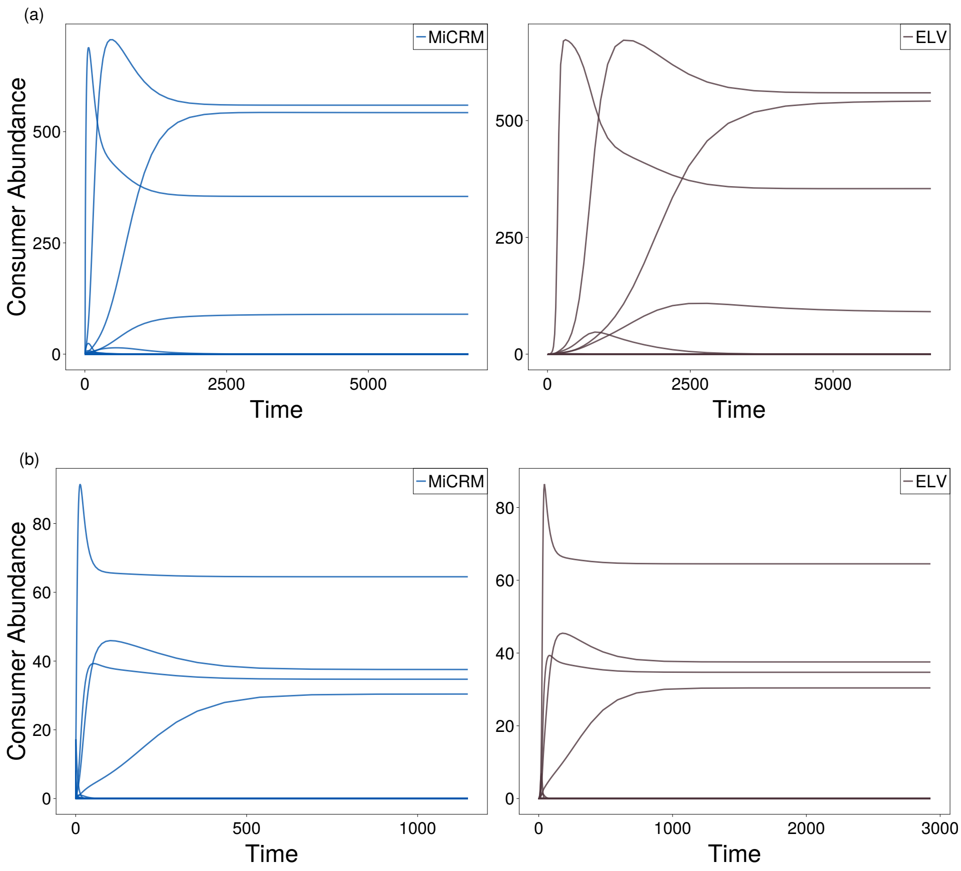


Figure S1: **The temporal dynamics of consumer abundance in MiCRM and ELV.** The simulations are run without thermal metabolic constraints, showing the trajectory of 100 species ($N = 100$) competing for 50 resource types ($M = 50$) at 0 °C (a) and 30 °C (b). Note that at higher temperatures, both systems reach steady states (exponentially) faster than at lower temperatures.

#### S2 The temperature-dependence of intra- and inter-specific interaction coefficients

Here, we explain three key results: i) Why the temperature-dependence of the mean effective intraspecific ($\alpha_{ii}$) and interspecific ($\alpha_{i\neq j}$) interaction coefficients differ (Fig. 3); ii) Why the effective pairwise species interactions are more thermally sensitive than their underlying metabolic traits (Fig. 2); iii) Why the ratios between interspecific and intraspecific interactions respond unimodally to temperature (Fig. 3d).

To this end, we will first derive and analyse the temperature dependence of effective coefficients of the simplest system that can provide the necessary insights: a 2 consumer-1 resource system without leakage (Eq. 8).

**(I) Why the temperature-dependence of effective intraspecific and interspecific effective interaction coefficients differ**

We start with the quasi-steady-state condition for the resource abundance $\hat{R}_a(\mathbf{C})$ in this simplified system. Since there is no leakage, the condition is:

$$\rho_a(\hat{R}_a) - \sum_i C_i u_{ia} \hat{R}_a = 0.$$

To find the interaction coefficients, we calculate the sensitivity of equilibrium resource abundance ($R_a = R_a^*$) to changes in consumer biomass. First we apply the chain rule following sec. S1 (Eqn. S4):

$$\frac{\partial \rho_a}{\partial \hat{R}_a} \cdot \frac{\partial \hat{R}_a}{\partial C_j} - \left( u_{ja} R_a^* + \sum_i C_i^* u_{ia} \frac{\partial \hat{R}_a}{\partial C_j} \right) = 0.$$

We derive further equations assuming the scenario where $\rho_a$ is the constant external resource supply of resource type $a$. In other scenarios, such as leaching ($\frac{\partial \rho_a}{\partial \hat{R}_a} = -\omega_a$) or chemostat ($\frac{\partial \rho_a}{\partial \hat{R}_a} = -\chi_a$) resource environments, the derivation is identical except for an addition of a constant term ($-\omega_a$ or $-\chi_a$). As for constant resource supply, $\frac{\partial \rho_a}{\partial \hat{R}_a} = 0$. We obtain the impact of species $j$ on equilibrium resource abundance of $a$:

$$\frac{\partial \hat{R}_a}{\partial C_j} = -\frac{u_{ja} R_a^*}{\sum_i C_i^* u_{ia}}. \tag{S10}$$

Substituting this derivative into the general definition of the interaction coefficient (Eq. S7) and noting that $l = 0$:

$$\begin{aligned} \alpha_{ij} &= u_{ia} \frac{\partial \hat{R}_a}{\partial C_j} \\ &= -u_{ia} \cdot \frac{u_{ja} R_a^*}{\sum_i C_i^* u_{ia}}. \end{aligned} \tag{S11}$$

From this, we obtain the explicit expressions for the intraspecific ($\alpha_{ii}$) and interspecific ($\alpha_{ij}$) coefficients:

$$\alpha_{ii} = -\frac{u_{ia}^2 R_a^*}{u_{ia} C_i^* + u_{ja} C_j^*}, \tag{S12}$$

$$\alpha_{ij} = -\frac{u_{ia} u_{ja} R_a^*}{u_{ia} C_i^* + u_{ja} C_j^*}. \tag{S13}$$

Note that here the interaction matrix $\mathbf{A}$ is symmetric ($\alpha_{ij} = \alpha_{ji}$) since we are considering the system without leakage function (i.e., without cross-feeding).

Consumer species here are interacting through just one resource type, and the temperature dependence of resource uptake is species-specific. We incorporate the Boltzmann–Arrhenius Equation to approximate the exponential subset of TPCs before high-temperature deactivation takes effect, with $\Delta T = -k(\frac{1}{T} - \frac{1}{T_r})$ (where k is the Boltzmann constant and $T_r$ is the reference temperature; see Section S3). Then the uptake of resource by each species can be written as:

$$u_{ia} = B_{u_i} e^{E_{u_i} \Delta T},$$
$$u_{ja} = B_{u_j} e^{E_{u_j} \Delta T}.$$

Here, the $B_u$s and $E_u$s are the normalization constants and activation energies of species-level uptake rates.

We incorporate the temperature dependence of resource uptake rates into Eq. S12 and Eq. S13, yielding:

$$\alpha_{ii} = -\frac{B_{u_i}^2 \cdot e^{2E_{u_i}\Delta T} R_a^*}{B_{u_i} e^{E_{u_i}\Delta T} C_i^* + B_{u_j} e^{E_{u_j}\Delta T} C_j^*}, \tag{S14}$$

$$\alpha_{ij} = -\frac{B_{u_i} B_{u_j} \cdot e^{(E_{u_i}+E_{u_j})\Delta T} R_a^*}{B_{u_i} e^{E_{u_i}\Delta T} C_i^* + B_{u_j} e^{E_{u_j}\Delta T} C_j^*}. \tag{S15}$$

Thus, here in the simplified one-resource model with no leakage, $\alpha_{ii}$ scales with the *square* of species $i$'s uptake rate, while $\alpha_{ij}$ scales with the *product* of the uptake rates of $i$ and $j$.

We can analyze the expected values of the interaction coefficients by first approximating them as:

$$-\alpha_{ii} \propto \frac{u_i^2}{D},$$
$$-\alpha_{ij} \propto \frac{u_i u_j}{D},$$

where $D$ is the common denominator term (equilibrium resource abundance and summed uptake).

As described in Methods section 2.4, both $u_i(T)$ and $u_j(T)$ follow a thermal performance curve (TPC), such as an Arrhenius or Sharpe–Schoolfield form, and $u_i$ and $u_j$ are random variables representing uptake rates drawn from the distribution of TPCs across species. The normalization constant $B_u$ and the activation energies $E_u$ of species-level resource uptake rates ($u_i$ and $u_j$) were drawn from the same distribution, that is $ln(B_u) \sim \mathcal{N}(\mu_{ln(B_u)}, \sigma^2_{ln(B_u)})$ and $E_u \sim \mathcal{N}(\mu_{E_u}, \sigma^2_{E_u})$.

Using Jensen's inequality, we can formally compare the expected values of these interactions.

For intraspecific interactions, the expected value of the square is related to the square of the expected value by the variance of the uptake rate, Var$[u]$:

$$\mathbb{E}[u_i^2] = \mathbb{E}[u^2] = (\mathbb{E}[u])^2 + \mathrm{Var}[u].$$

Since variance is non-negative, this implies $\mathbb{E}[u_i^2] \geq (\mathbb{E}[u])^2$, with equality only if uptake rates are identical across all species.

Conversely, for interspecific interactions, since the traits of species $i$ and $j$ are independent, the expected value of the product is simply the product of the expected values:

$$\mathbb{E}[u_i u_j] \;=\; \mathbb{E}[u_i] \cdot \mathbb{E}[u_j] \;=\; \big(\mathbb{E}[u]\big)^2.$$

Comparing these two results reveals that:

$$\mathbb{E}[-\alpha_{ii}] \geq \mathbb{E}[-\alpha_{ij}].$$

This demonstrates that, on average, intraspecific interaction coefficients will increase faster than interspecific interactions with warming.

This is especially true since that, as temperature increases, the variance of uptake rates across species typically grows. This is because the underlying parameters ($B_u$ and $E_u$) are distributed exponentially; small differences in activation energy $E_u$ lead to increasingly large divergences in uptake rates $u$ at higher temperatures. Consequently, since

$$\mathbb{E}[u_i^2] - \mathbb{E}[u_i u_j] \; = \; \mathrm{Var}[u],$$

the gap between $\mathbb{E}[\alpha_{ii}]$ and $\mathbb{E}[\alpha_{ij}]$ widens with temperature.

This mathematical property leads to two key consequences for the effective Lotka-Volterra (ELV) coefficients:

i. $\mathbb{E}[\alpha_{ii}]$ grows disproportionately fast with temperature. Because $\alpha_{ii}$ involves the square of a random variable (the trait $u_i$), its expected value includes extra exponential factors from the growing variance of species' uptake rates.

ii. $\mathbb{E}[\alpha_{ij}]$, by contrast, grows more slowly. Because it depends on the product of independent uptake rates, and is therefore less sensitive to the widening variance of the underlying metabolic distribution.

This analytical insight aligns with our simulation results (Fig. 3b), where the mean intraspecific interaction strength increases faster with temperature than the mean interspecific interaction strength.

#### (II) Why effective pairwise species interactions are more thermally sensitive than their underlying metabolic traits

To compare the temperature performances of the mean values of $\alpha_{ii}$ and $\alpha_{i\neq j}$ more precisely, following equations S14 and S15, we write out the equations of $\mathbb{E}[\alpha_{ii}]$ and $\mathbb{E}[\alpha_{ij}]$ as follows:

$$\begin{aligned}\mathbb{E}[\alpha_{ii}] &= \left( \frac{R_a^* \cdot \mathbb{E}[B_{u_i}^2 e^{2E_{u_i}\Delta T}]}{\mathbb{E}[B_{u_i} e^{E_{u_i}\Delta T} C_i^* + B_{u_j} e^{E_{u_j}\Delta T} C_j^*]} \right) \cdot \phi_{\alpha_{ii}} \\ &\approx \frac{R_a^*}{C_i^* + C_j^*} \cdot \frac{e^{2\mu_{\ln(B_u)} + 2\sigma^2_{\ln(B_u)} + 2\mu_{E_u}\Delta T + 2\sigma^2_{E_u}\Delta T^2}}{e^{\mu_{\ln(B_u)} + \mu_{E_u}\Delta T + \frac{1}{2}\sigma^2_{\ln(B_u)} + \frac{1}{2}\sigma^2_{E_u}\Delta T^2}} \\ &= \frac{R_a^*}{C_i^* + C_j^*} \cdot e^{\mu_{\ln(B_u)} + \frac{3}{2}\sigma^2_{\ln(B_u)} + \mu_{E_u}\Delta T + \frac{3}{2}\sigma^2_{E_u}\Delta T^2},\end{aligned}$$

and

$$\begin{aligned}\mathbb{E}[\alpha_{ij}] &= \left( \frac{R_a^* \cdot \mathbb{E}[B_{u_i} B_{u_j} e^{(E_{u_i} + E_{u_j})\Delta T}]}{\mathbb{E}[B_{u_i} e^{E_{u_i}\Delta T} C_i^* + B_{u_j} e^{E_{u_j}\Delta T} C_j^*]} \right) \cdot \phi_{\alpha_{ij}} \\ &\approx \frac{R_a^*}{C_i^* + C_j^*} \cdot \frac{e^{2\mu_{\ln(B_u)} + \sigma^2_{\ln(B_u)} + 2\mu_{E_u}\Delta T + \sigma^2_{E_u}\Delta T^2}}{e^{\mu_{\ln(B_u)} + \mu_{E_u}\Delta T + \frac{1}{2}\sigma^2_{\ln(B_u)} + \frac{1}{2}\sigma^2_{E_u}\Delta T^2}} \\ &= \frac{R_a^*}{C_i^* + C_j^*} \cdot e^{\mu_{\ln(B_u)} + \frac{1}{2}\sigma^2_{\ln(B_u)} + \mu_{E_u}\Delta T + \frac{1}{2}\sigma^2_{E_u}\Delta T^2}.\end{aligned}$$

Here $\phi_{\alpha_{ii}}$ and $\phi_{\alpha_{ij}}$ are correction factors accounting for the covariance between the numerator and denominator in each interaction term, and the effects of the variance of the denominator (which is identical for both $\alpha_{ii}$ and $\alpha_{ij}$). Since both the covariance and variance terms of the numerator and denominator scale similarly with temperature, as the normalisation constants $B_u$ and activation energies $E_u$ of both species $i$ and $j$ are drawn from the same trait distributions, the temperature dependence of $\phi_{\alpha_{ii}}$ and $\phi_{\alpha_{ij}}$ is not expected to differ significantly. Therefore, the observed faster increase of $\mathbb{E}[\alpha_{ii}]$ with temperature compared to $\mathbb{E}[\alpha_{ij}]$ can be attributed to the stronger propagation of trait variance in the exponential terms of their expected values.

To compare the temperature performance curves of $\alpha_{ii}$ and $\alpha_{ij}$, we first take the natural logarithm of $-\alpha_{ii}$ (Eqn. S14):

$$ln(-\alpha_{ii}) = ln(B_{u_i}^2 R_a^*) + 2E_{u_i}\Delta T - ln(B_{u_i}e^{E_{u_i}\Delta T}C_i^* + B_{u_j}e^{E_{u_j}\Delta T}C_j^*).$$

Then compute the first-order Taylor expansion of $ln(-\alpha_{ii})$ around $\Delta T = 0$:

$$\ln(-\alpha_{ii}) \approx ln\left(\frac{B_{u_i}^2 R_a^*}{B_{u_i}C_i^* + B_{uj}C_j^*}\right) + \left(2E_{u_i} - \frac{B_{u_i}E_{u_i}C_i^* + B_{u_j}E_{u_j}C_j^*}{B_{u_i}C_i^* + B_{u_j}C_j^*}\right)\Delta T.$$

Here we show that the temperature performance of effective intra-specific interactions $\alpha_{ii}$ is Arrhenius-like within the OTR. The normalisation constant $B_{\alpha_{ii}}$ and activation energy $E_{\alpha_{ii}}$ are:

$$B_{\alpha_{ii}} = \frac{B_{u_i}^2 R_a^*}{B_{u_i}C_i^* + B_{uj}C_j^*}, \tag{S16}$$

$$E_{\alpha_{ii}} = 2E_{u_i} - \frac{B_{u_i}E_{u_i}C_i^* + B_{u_j}E_{u_j}C_j^*}{B_{u_i}C_i^* + B_{u_j}C_j^*}. \tag{S17}$$

Same way, we can derive the temperature performance equation of effective inter-specific interactions $\alpha_{ij}$ (Eqn. S15), where

$$ln(-\alpha_{ij}) = ln(B_{u_i}B_{u_j}R_a^*) + (E_{u_i} + E_{u_j})\Delta T - ln(B_{u_i}e^{E_{u_i}\Delta T}C_i^* + B_{u_j}e^{E_{u_j}\Delta T}C_j^*).$$

Computing the first-order Taylor expansion of $ln(-\alpha_{ij})$ around $\Delta T = 0$:

$$\ln(-\alpha_{ij}) \approx ln\left(\frac{B_{u_i}B_{u_j}R_a^*}{B_{u_i}C_i^* + B_{uj}C_j^*}\right) + \left(E_{u_i} + E_{u_j} - \frac{B_{u_i}E_{u_i}C_i^* + B_{u_j}E_{u_j}C_j^*}{B_{u_i}C_i^* + B_{u_j}C_j^*}\right)\Delta T.$$

The normalisation constant $B_{\alpha_{ij}}$ and activation energy $E_{\alpha_{ij}}$ of the temperature performance of $\alpha_{ij}$ are:

$$B_{\alpha_{ij}} = \frac{B_{u_i}B_{u_j}R_a^*}{B_{u_i}C_i^* + B_{uj}C_j^*}, \tag{S18}$$

$$E_{\alpha_{ij}} = E_{u_i} + E_{u_j} - \frac{B_{u_i}E_{u_i}C_i^* + B_{u_j}E_{u_j}C_j^*}{B_{u_i}C_i^* + B_{u_j}C_j^*}. \tag{S19}$$

Here, in both Eqn. S17 and S19, the second terms are identical. Therefore we denote $X = \frac{B_{u_i}E_{u_i}C_i^* + B_{u_j}E_{u_j}C_j^*}{B_{u_i}C_i^* + B_{u_j}C_j^*}$. Then the activation energies simplify to:

$$E_{\alpha_{ii}} = 2E_{u_i} - X,$$
$$E_{\alpha_{ij}} = E_{u_i} + E_{u_j} - X.$$

Hence, the activation energies of both interspecific interactions $E_{\alpha_{ij}}$ and intraspecific interactions $E_{\alpha_{ii}}$ are equal, where $\mathbb{E}[E_{\alpha_{ii}}] = \mathbb{E}[E_{\alpha_{ij}}] = 2\mu_{E_u} - \mathbb{E}[X]$, as shown in Fig. S5.

In realistic scenarios where there exists a negative correlation between the normalisation constants and activation energies of species-level uptake rates (i.e. between $B_u$ and $E_u$) due to the thermal-generalist and specialist trade-off (see detailed description in Sec. S4), we have

$$\mathbb{E}[B_u \cdot E_u] = \mathbb{E}[B_u] \cdot \mathbb{E}[E_u] + \rho_{B_u,E_u} \cdot \sigma_{B_u} \cdot \sigma_{E_u}, \text{ with } \rho_{B_u,E_u} < 0.$$

This implies that

$$\mathbb{E}[X] < \frac{(C_i^* + C_j^*) \cdot \mathbb{E}[B_u] \cdot \mu_{E_u}}{(C_i^* + C_j^*) \cdot \mathbb{E}[B_u]} = \mu_{E_u}.$$

Therefore, the effective activation energies of species interactions satisfy $\mathbb{E}[E_\alpha] > \mu_{E_u}$. Since the mean of activation energies of species uptake rates $E_u$ are known to exceed those of respiration rates $E_m$ (see Methods section 2.4), it follows that $\mathbb{E}[E_\alpha] > \mu_{E_u} > \mu_{E_m}$.

We thus conclude that effective pairwise species interactions are typically more thermally sensitive than their underlying metabolic traits, including both uptake and respiration rates (Fig. 2).

**(III) Why the ratios between interspecific and intraspecific interactions respond unimodally to temperature**

Following Eq. S14 and Eq. S15, the ratio between the interspecific and intraspecific interaction coefficients can be expressed as:

$$\frac{\alpha_{ij}}{\alpha_{ii}} = \frac{B_{u_j}}{B_{u_i}} e^{(E_{u_j} - E_{u_i})\Delta T}.$$

Taking the logarithm gives:

$$ln(\frac{\alpha_{ij}}{\alpha_{ii}}) = \left(lnB_{u_j} - lnB_{u_i}\right) + (E_{u_j} - E_{u_i})\Delta T.$$

Then, the expected values of this ratio is:

$$\mathbb{E}[ln(\frac{\alpha_{ij}}{\alpha_{ii}})] = \mathbb{E}[lnB_{u_j} - lnB_{u_i}] + \mathbb{E}[(E_{u_j} - E_{u_i})\Delta T] + \frac{1}{2} \cdot \left(2\sigma^2_{lnB_u} + 2\sigma^2_{E_u}\Delta T^2 + 4\rho_{lnB_u,E_u}\sigma_{lnB}\sigma_E \Delta T\right).$$

Because both $i$ and $j$ are drawn independently from the same distribution of $\ln B_u$ and $E_u$, we have $\mathbb{E}[\ln B_{u_j}] = \mathbb{E}[\ln B_{u_i}]$ and $\mathbb{E}[E_{u_j}] = \mathbb{E}[E_{u_i}]$. Therefore,

$$\mathbb{E}[lnB_{u_j} - lnB_{u_i}] = 0,$$
$$\mathbb{E}[(E_{u_j} - E_{u_i}) = 0.$$

This leaves only the variance and covariance terms that determine how the ratio changes with temperature:

$$\mathbb{E}[ln(\frac{\alpha_{ij}}{\alpha_{ii}})] = \sigma^2_{lnB_u} + \sigma^2_{E_u}\Delta T^2 + 2\rho_{lnB_u,E_u}\sigma_{lnB}\sigma_{E_u}\Delta T.$$

Taking the derivative with respect to $\Delta T$ and setting it to zero gives:

$$2\sigma^2_{E_u}\Delta T + 2\rho_{lnB_u,E_u}\sigma_{lnB}\sigma_{E_u} = 0.$$

Therefore,

$$\Delta T = -\rho_{lnB_u,E_u}\frac{\sigma_{lnB}}{\sigma_{E_u}}$$

Since $\rho_{lnB_u,E_u} < 0$, we have $\Delta T > 0$, meaning that the ratio between interspecific and intraspecific interaction coefficients $\alpha_{ij}/\alpha_{ii}$ responds unimodally to temperature, with a minimum at an intermediate temperature above the reference $T_r$.

#### S3 Thermal performance curve parameterisations and diffusion limitation

Here, we show that our empirically derived parameterizations (Table S1) of the framework implicitly capture the effects of diffusion limitation.

Table S2: **Parameters for the temperature dependencies of uptake and respiration.**

| Symbols | Definition | Value | Units |
|---|---|---|---|
| $\mu_{\ln(B_{0,u})}$ | Mean normalization constant for uptake rate | $-0.81$ | - |
| $\mu_{\ln(B_{0,m})}$ | Mean normalization constant for respiration rate | $-1.50$ | - |
| $\sigma^2_{\ln(B_{u,0})}$ | Variance of normalization constant for uptake rates | 0.17 | - |
| $\sigma^2_{\ln(B_{m,0})}$ | Variance of normalization constant for respiration rates | 0.17 | - |
| $\mu_{E_u}$ | Mean activation energy for uptake rate | 0.82 | eV |
| $\mu_{E_m}$ | Mean activation energy for respiration rate | 0.57 | eV |
| $\sigma^2_{E_u}$ | Variance of activation energy for uptake rate | 0.14 | $eV^2$ |
| $\sigma^2_{E_m}$ | Variance of activation energy for respiration rate | 0.14 | $eV^2$ |
| $\mu_{T_{pk,u}}$ | Mean peak temperature for uptake rate | 35 | $^o$C |
| $\mu_{T_{pk,m}}$ | Mean peak temperature for respiration rate | 38 | $^o$C |
| $\sigma^2_{T_{pk,u}}$ | Variance of peak temperature for uptake rate | 5 | $^o$C$^2$ |
| $\sigma^2_{T_{pk,m}}$ | Variance of peak temperature for respiration rate | 5 | $^o$C$^2$ |
| $k$ | Boltzmann constant | $8.62 * 10^{-5}$ | eV/K |
| $T$ | Model temperature | $0 \sim 30$ (38) | $^o$C |
| $T_r$ | Reference temperature | 10 | $^o$C |
| $E_{d,u}$ | Deactivation energy for uptake rate | 3.5 | eV |
| $E_{d,m}$ | Deactivation energy for respiration rate | 3.5 | eV |

We extracted TPC parameterizations from empirical data by fitting the bacterial dataset (Methods) to the Sharpe-Schoolfield TPC equation using non-linear least squares fitting with the R package `rTPC` (Padfield et al. 2021). Because very few species in the dataset were psychrophiles (extreme cold-tolerant), we chose 10°C as the reference temperature ($T_r$) to minimize the extrapolation error in the normalized $B_0$.

Fitting the Sharpe-Schoolfield equation to experimentally measured growth-rate and respiration data implicitly includes diffusion limitation. The experimental measures of these two traits necessarily include diffusion limitation because nutrient molecules must diffuse to the cell surface before uptake, and metabolites must diffuse through the cytoplasm before enzyme-mediated reactions occur (Kiørboe 2009). Maximum nutrient uptake rates scale approximately with cell surface area and the abundance of transporters, and half-saturation constants scale with cell radius, reflecting diffusion-limited transport (Andersen & Visser 2023, Fenchel 1974, Moloney & Field 1991). Similarly, studies of bacterial proteome dynamics show that intracellular protein diffusion strongly constrains metabolic rates and decreases at high temperatures, further embedding diffusion limitation into empirically-observed TPCs (Schavemaker et al. 2018, Di Bari et al. 2023).

Therefore, the normalisation constant $B_0$ in our parameterisations is also expected to scale with cell size, partly due to diffusion limitation: smaller cells have a proportionally larger surface area relative to volume and thus exhibit higher uptake affinities. For example, in phytoplankton, maximum nutrient uptake rates scale with cell surface area (size exponent $\approx 2/3$), whereas half-saturation constants scale with cell radius (exponent $\approx 1/3$), as predicted by mass-transfer theory and consistent with early work on protozoa and phytoplankton showing that specific nitrate uptake scales with cell volume to a power of approximately $-1/3$ (Fenchel 1974, Moloney

& Field 1991, Tang & Peters 1995). Because diffusion to a spherical cell surface is proportional to the cell radius $R$, the steady-state diffusive flux of nutrients is $F_{\mathrm{diff}} = 4\pi D R \Delta C$, where $D$ is diffusivity and $\Delta C$ the concentration gradient. Thus, doubling $R$ doubles the total diffusive flux, but reduces the per-biomass uptake rate as $R^{-1}$ or $V^{-1/3}$, implying that smaller cells achieve higher normalisation constants under diffusion limitation, with a log-normal-like distribution (Fig. S2 a).

Thus, our empirically derived TPCs for $u$ (approximated from $r_{\max}$) and $m$ implicitly capture both enzyme-kinetic and diffusion-limited processes. The observed variation in our estimated activation energies $E$ and $B_0$ is consistent with this, as we show below.

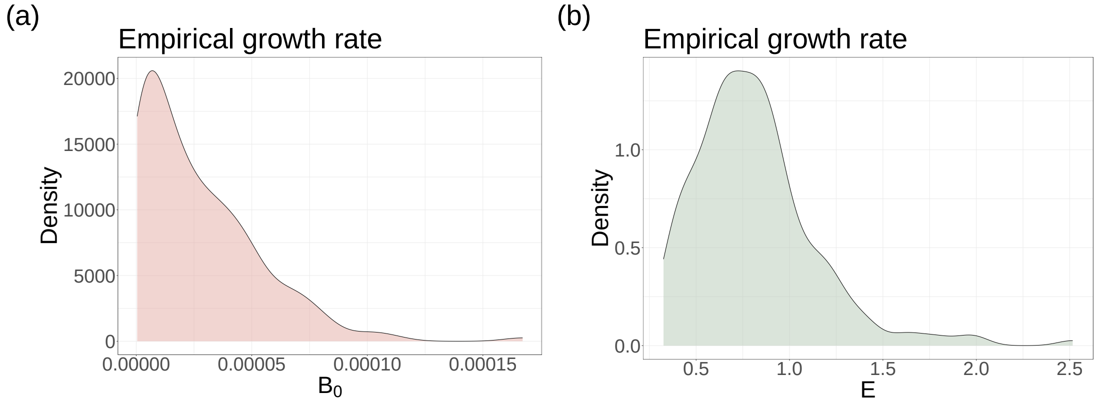


Figure S2: **The distribution of the normalisation constant ($B_0$; a) and the activation energy ($E$; b) of growth rate from empirical data.**

Second, the temperature dependence of diffusion is implicit in the estimates of $E$ obtained by fitting the Sharpe-Schoolfield to the growth-rate data. The molecular diffusion coefficient $D$ of small solutes in water obeys an Arrhenius-type relation

$$D(T) = D_0 e^{-E_{diff}/(kT)}, \tag{S20}$$

where $D_0$ is a pre-exponential factor and $E_{diff}$ is the activation energy associated with diffusive motion (Mills 1973, Piskulich et al. 2017). Measurements of self-diffusion in water indicate $E_{diff} \approx 18\,\mathrm{kJ\,mol^{-1}}$ (Mills 1973). When nutrient uptake is diffusion-limited, the overall uptake rate $u(T)$ is proportional to $D(T)$, leading to an effective temperature dependence

$$u(T) \propto D(T) u_{\mathrm{enz}}(T), \tag{S21}$$

where $u_{\mathrm{enz}}(T)$ is the enzyme-kinetic contribution described by the Sharpe–Schoolfield TPC model. Similarly, intracellular diffusion of metabolites influences the apparent respiration rate $m(T)$: biophysical studies of protein diffusion in *Escherichia coli* show that proteome diffusion exhibits a linear Stokes–Einstein dependence on temperature at sublethal temperatures and slows down dramatically near the cell's death temperature due to protein unfolding (Bellotto et al. 2022, Di Bari et al. 2023, Śmigiel et al. 2022). Thus, both extracellular and intracellular diffusion exhibit Arrhenius-like temperature dependence (Weber et al. 2012, Bag et al. 2014) and can be incorporated into the effective activation energies of uptake and respiration. Therefore, replacing $\mathrm{e}^{-E/(kT)}$ in the Sharpe-Schoolfield equation with the product $\mathrm{e}^{-(E+E_{diff})/(kT)}$ yields a modified TPC in which the apparent activation energy $E_{\mathrm{app}} = E + E_{diff}$, that includes the contribution from diffusion. Because $E_{diff} \ll E$ for most metabolic reactions, this modification increases the slope of the Arrhenius plot by only a modest amount as we now explain.

Estimates of activation energies for microbial respiration and resource uptake span the range 0.38–0.65 eV (Corkrey et al. 2012, Pirt 1975). In contrast, the activation energy of molecular diffusion in water is approximately 0.19 eV (Mills 1973). Assuming that effective uptake (and respiration) rates are the product of an enzyme-kinetic rate with activation energy $E$ and a diffusion term with activation energy $E_{diff}$, the apparent activation energy is $E_{\text{app}} = E + E_{diff}$. Table S3 summarises the fractional contribution of diffusion limitation to the estimated apparent activation energy across representative scenarios. When $E = 0.65$ eV, diffusion accounts for $E_{diff}/(E + E_{diff}) \approx 23\%$. When $E = 0.40$ eV, diffusion contributes roughly 33%. Even in the most diffusion-limited scenarios, the diffusion component contributes less than half of the apparent activation energy; thus, our qualitative results—which depend on the relative thermal sensitivities of uptake and respiration—are robust across a broad range of $E_{diff}$ values.

Varying the diffusion activation energy $E_{diff}$ within plausible bounds (e.g. 0–0.25 eV) changes the apparent activation energies by at most a few tenths of an eV (Table S3), well within the range explored in our analyses. Consequently, the key predictions of our framework—the higher thermal sensitivity of effective interactions relative to individual metabolic traits and the temperature dependence of self-regulation versus interspecific interactions, already accounting for diffusion limitation.

Table S3: Illustrative contribution of diffusion limitation to the apparent activation energy of uptake or respiration. We assume $E_{diff} = 0.196$ eV for diffusion and vary the enzyme-kinetic activation energy $E$ of the underlying metabolic process. The diffusion fraction is $E_{diff}/(E + E_{diff})$.

| Enzyme activation energy $E$ (eV) | Apparent $E_{\text{app}}$ (eV) | Diffusion fraction (%) |
|---|---|---|
| 0.65 | 0.846 | 23 |
| 0.50 | 0.696 | 28 |
| 0.40 | 0.596 | 33 |
| 0.30 | 0.496 | 40 |

In summary, our parameterisations use empirical TPCs that already encapsulate diffusion, and our theoretical results about the temperature dependence of microbial interactions are robust to alternative TPC models and realistic variations in diffusion limitation. Indeed, the Sharpe–Schoolfield model is a convenient but not exclusive choice for describing temperature-dependent microbial interactions, and any unimodal TPC can replace it within our framework.

#### S4 Thermal generalist-specialist tradeoff as an additional metabolic constraint

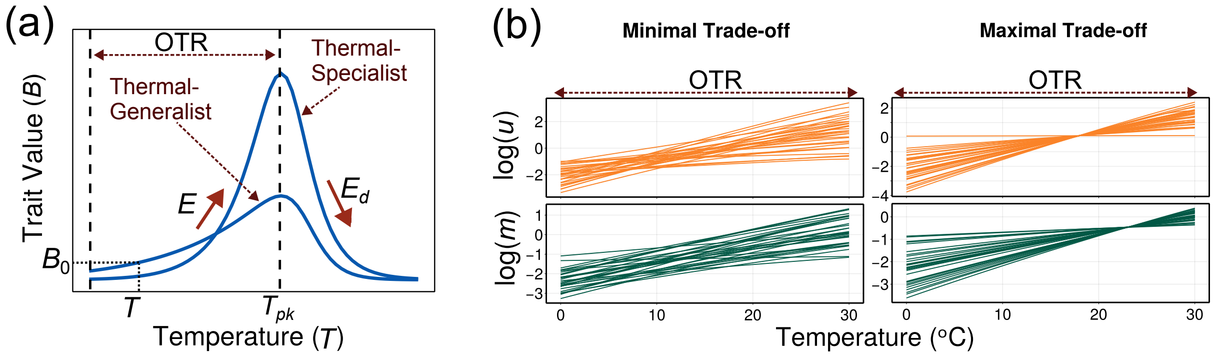


Figure S3: **Modelling thermal generalists, specialists, and the generalist-specialist tradeoff.**

In ectotherms, acclimatization or adaptation to one environmental temperature generally comes at the cost of performance at another (Mongold et al. 1996, Angilletta et al. 2002, Ketola & Saarinen 2015). Therefore, TPCs of performance traits (such as $u$ and $m$) across ectotherms, including microbes, typically exhibit a (thermal) generalist-specialist trade-off (Fig. S3). That is, TPCs of thermal generalists show moderate levels of performance across a wide range of temperatures, but do not achieve exceptionally high performance at any specific temperature, while those of thermal specialists show high performance within a narrow temperature range but exhibit steep declines in performance outside this range (Huey & Hertz 1984, Gilchrist 1995). By fixing the reference temperature $T_r$ at the lower end of the operational temperature range, this thermal generalist-specialist trade-off can be quantified as a negative correlation between activation energy $E$ and normalisation constant $B_0$ ($\rho_{\ln(B_0),E}$; Fig. 1c; Eqn. 9; (Kontopoulos, van Sebille, Lange, Yvon-Durocher, Barraclough & Pawar 2020, Kontopoulos, Smith, Barraclough & Pawar 2020)).

We assume that in any local community assembly dynamic, the value of the thermal generalist-specialist tradeoff $\rho_{\ln(B_0),E}$ varies across species that encounter each other. That is, both thermal generalists and specialists are expected to encounter each other in a local environment (or patch), for two reasons. First, those different species will have inherent differences in the strength of their trade-off ($\rho_{\ln(B_0),E}$) due to differences in their metabolic architecture (e.g., they may represent different phylogenetic groups or metabolic strategies; (Smith et al. 2019, Kontopoulos, Smith, Barraclough & Pawar 2020)). Second, different species in a local environment will likely have adapted to different ecological contexts, especially if the immigration pool covers a larger geographical area. Moreover, the generation times of species within any microbial local assemblage vary across orders of magnitude (from minutes to months or longer) (Gibson et al. 2018), and therefore they experience and adapt to effectively different regimes of environmental temperature fluctuations (Gilchrist 1995, Kontopoulos, van Sebille, Lange, Yvon-Durocher, Barraclough & Pawar 2020).

Therefore, the correlation between the normalization constant ($\ln(B_0)$) and the activation energy ($E$) within a microbial community is expected to vary depending on its evolutionary and ecological contexts. In theory, this trade-off could vary between two extremes. At one end (minimal trade-off, $\rho_{\ln(B_0),E} \approx 0$), both thermal generalists and specialists exist equally in a local patch. This scenario may arise when the geographical region from which the local community is assembled is large enough so that the species are drawn from multiple different thermal environments. Alternatively, it may arise when the variation in generation times is sufficiently high, allowing species to experience and adapt to frequent thermal fluctuations in their local environments (e.g., in higher-latitude regions). At the other end (maximal trade-off, $\rho_{\ln(B_0),E} \approx -1$), all species are thermal specialists. This scenario would arise when all species are preadapted to a constant environment with no thermal fluctuations, and have no intrinsic variation in the trade-off. Clearly, neither of these extreme scenarios is realistic. For example, even at lower geographical latitudes (the tropics), microbes experience thermal fluctuations at different timescales. Therefore, the trade-off levels of natural communities are expected to lie between these two extremes. Accordingly, in our simulations, we use an intermediate covariance value $\rho_{\ln(B_0),E} \approx -0.5$ to represent the realistic trade-off level between thermal generalists and specialists.

Finally, note that we assume that the value of $\rho_{B_0,E}$ is the same for $u$ and $m$. This is based on the fact that current empirical evidence, albeit limited, does not suggest any systematic difference in the correlation between these parameters across microbial trait types (Kontopoulos, Smith, Barraclough & Pawar 2020).

### S5 Thermal Sensitivity of Equilibrium Abundances

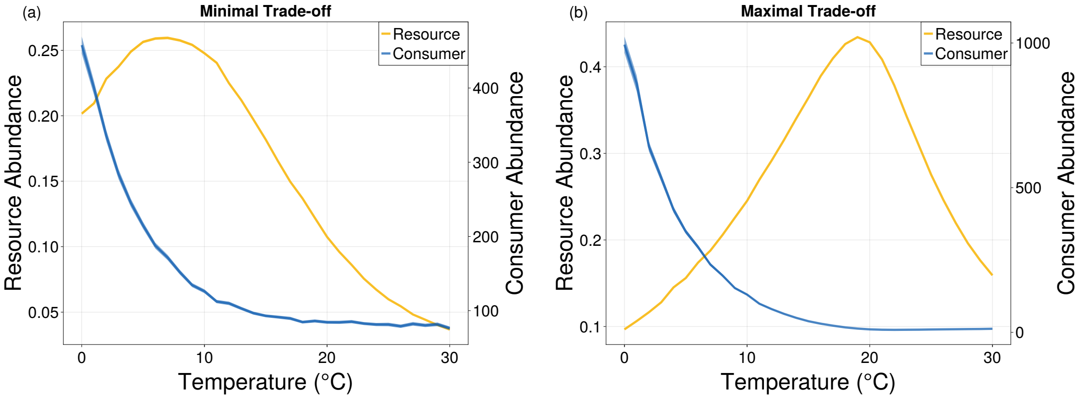


Figure S4: **Thermal responses of consumer and resource equilibrium abundance.** The mean equilibrium abundances of resources ($\bar{R}^*$; ±SE, yellow) and consumers ($\bar{C}^*$; ±SE, blue) are shown across the operational temperature range (OTR) for the two extreme cases described in Sec. S4: the minimal (a; $\rho_{\ln(B_0),E} = -0.5$) and maximal (b; $\rho_{\ln(B_0),E} \approx -1$) trade-off between thermal generalist-specialist. In both cases, mean consumer abundance follows a monotonic decline, while mean resource abundance follows a unimodal response peaking at intermediate temperatures. Higher trade-off levels (higher $\rho_{\ln(B_0),E}$) shift this peak to the right. Notably, both consumer and resource abundances decrease exponentially at relatively high temperatures, respiratory and resource uptake are still ascending.

#### S6 Temperature Dependence of Effective $\alpha_{ii}$ and $\alpha_{i\neq j}$

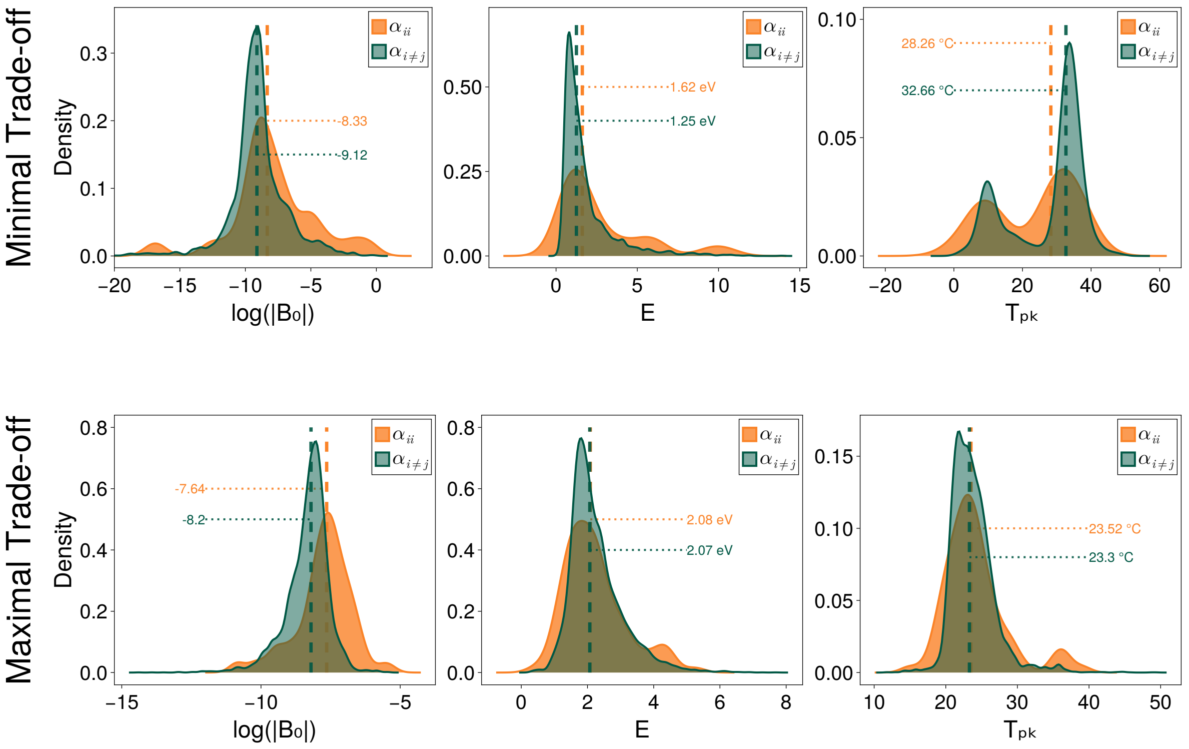


Figure S5: **The temperature dependence of effective intraspecific and interspecific interactions.** Intraspecific and interspecific interactions respond similarly to temperature, especially in the community with maximal trade-off, the main deviation lies in the difference between $B_0$.

#### S7 Robustness of our approach and its key predictions

To assess the generality of our framework, we examined whether the characteristics of the temperature dependence of effective interactions differ under different forms of resource supply and at different leakage levels. In addition to constant external input, we considered environments in which resources are lost from the system at a constant rate ($\rho_a = \rho_a^{in} - \omega_a R_a$), and chemostat environments (Tilman 1982, Marsland et al. 2019), in which resource input is described by $\rho_a = \chi_a(K_a - R_a)$, where $K_a$ and $\chi_a$ respectively denote the supply concentration and dilution rate of resource type $a$.

To align with classical consumer–resource models, we also examined the case of zero leakage, which corresponds to competitive consumer–resource systems without cross-feeding and is widely used beyond microbial systems. Across all resource regimes and leakage scenarios considered, the qualitative characteristics of the temperature dependence of effective interactions remain unchanged. Specifically, effective interactions exhibit unimodal thermal performance curves and are more thermally sensitive compared with their underlying metabolic traits (Fig. S6); interaction strengths increase with temperature across the operational temperature range, and the distributions of both intra- and interspecific interactions shift systematically with temperature (Fig. S7).

These results demonstrate that our main conclusions are not reliant on a particular form of resource supply, cross-feeding architecture, or leakage level, and support the generality of our framework across a wide range of environmentally relevant microbial settings. Analytical support for the invariance of these patterns under different resource regimes is provided in Sec. S2.

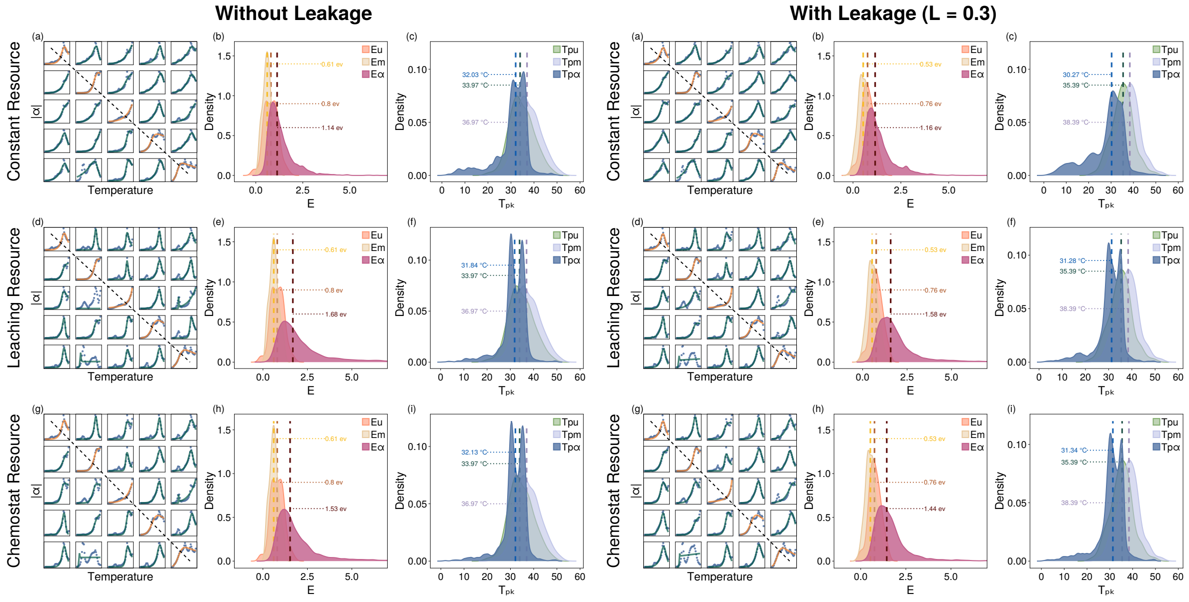


Figure S6: **The temperature performance curves (TPCs) of effective interactions under different resource input types, with and without leakage.**

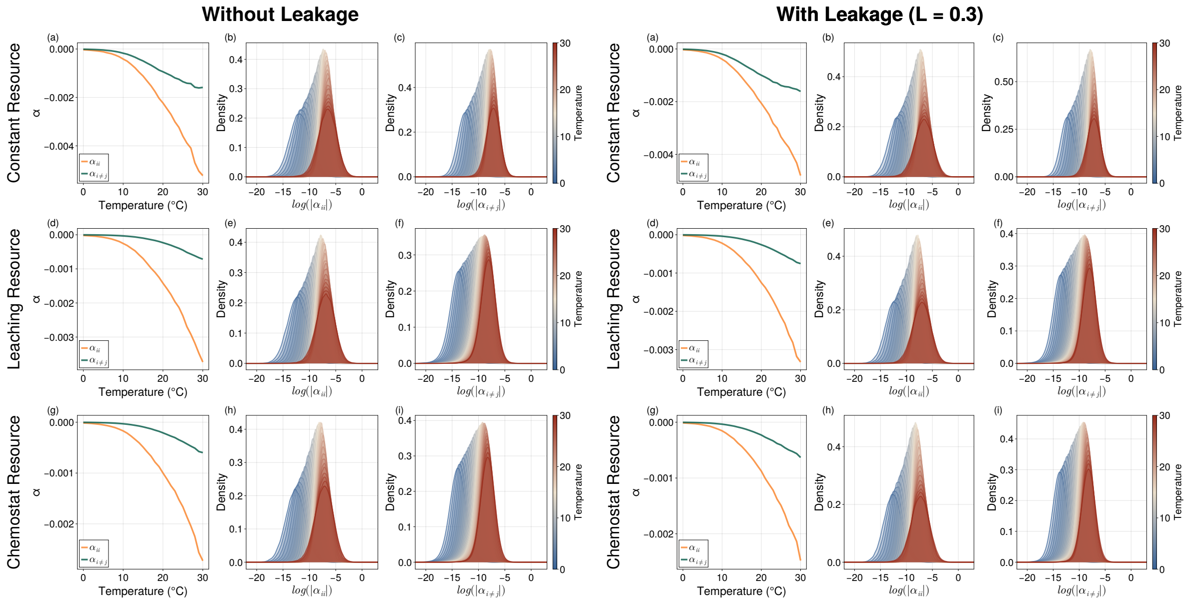


Figure S7: **The effect of temperature on effective interaction strengths under different resource input types, with and without leakage.**

#### S8 Effective $\alpha_{ii}$ and $\alpha_{i\neq j}$ without Thermal Metabolic Constraints

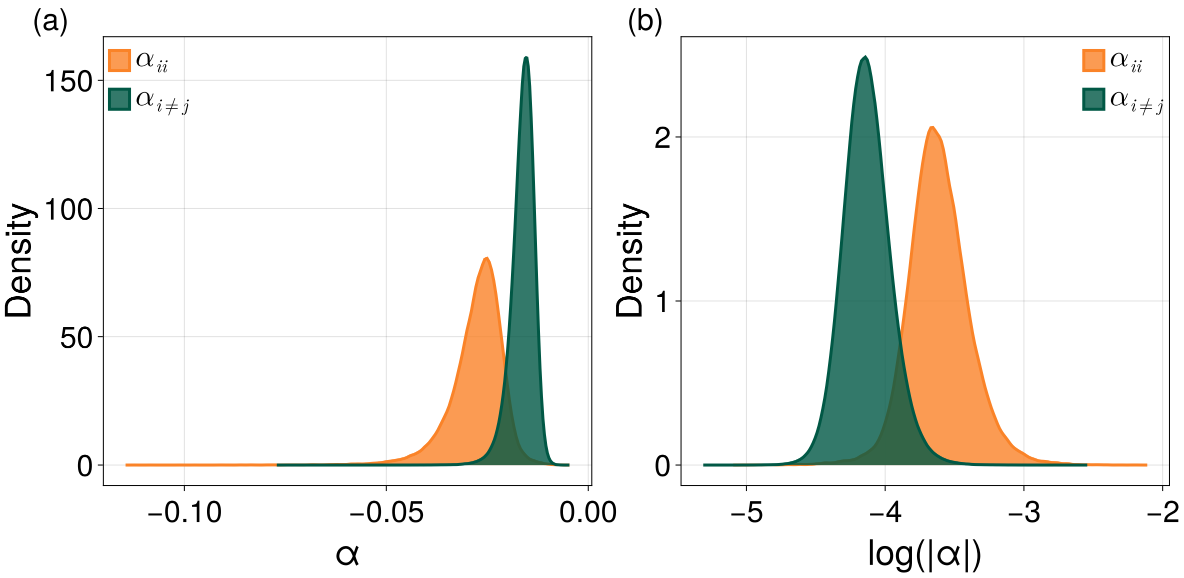


Figure S8: **The effective $\alpha_{ii}$ and $\alpha_{i\neq j}$ that emerge from consumer-resource dynamics are typically negative and follow a log-normal distribution.** The simulation was performed using normal distributions for both metabolic traits ($u$ and $m$) without thermal metabolic constraints ($\sigma \approx 0.1$).

#### S9 Temperature Dependence of Effective Interactions with Minimal Niche Overlap

In this case, one of the 50 resource types (the preferential resource type) is randomly assigned $\alpha = 100$, while the rest are assigned $\alpha = 1$ when generating the Dirichlet distribution. To achieve minimal niche overlap (high niche differentiation) in the community of 100 species competing for 50 resource types, each preferential resource type can only be shared by two randomly drawn species.

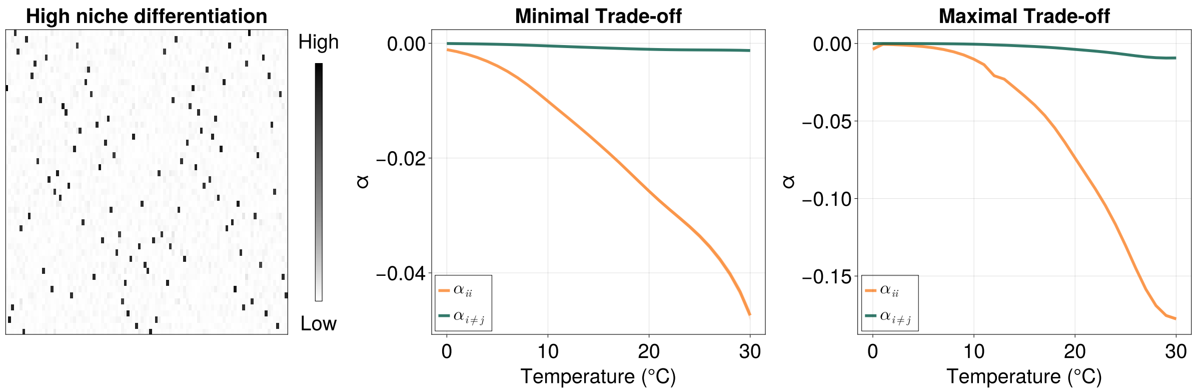


Figure S9: **Temperature dependence of effective interactions in communities with minimal niche overlap.** The heatmap shows the uptake matrix and the shape depicts the resource preferences of each species.

#### S10 Temperature Dependence of Interactions at Different Leakage Levels

To test the robustness of these results on the temperature dependencies of effective pairwise interactions on leakage levels, we run the simulations for a total of 999 assemblies across different leakage fractions from 0.1 to 0.7 (Fig. S10).

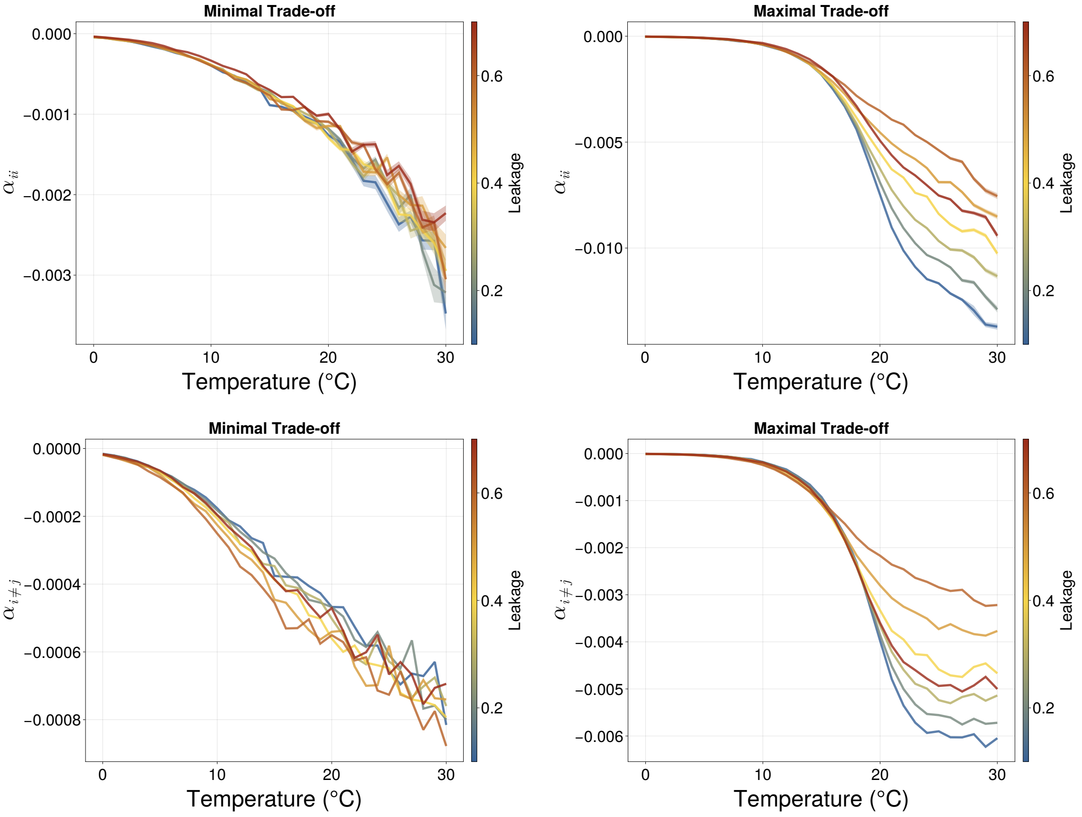


Figure S10: **Temperature dependence of effective interactions across different leakage levels.** The results qualitatively hold for all leakage levels from 0.1 to 0.7, where interaction strengths always increase with rising temperatures, where intraspecific interactions increase disproportionally faster.

#### S11 The TPCs of effective interactions within an example community

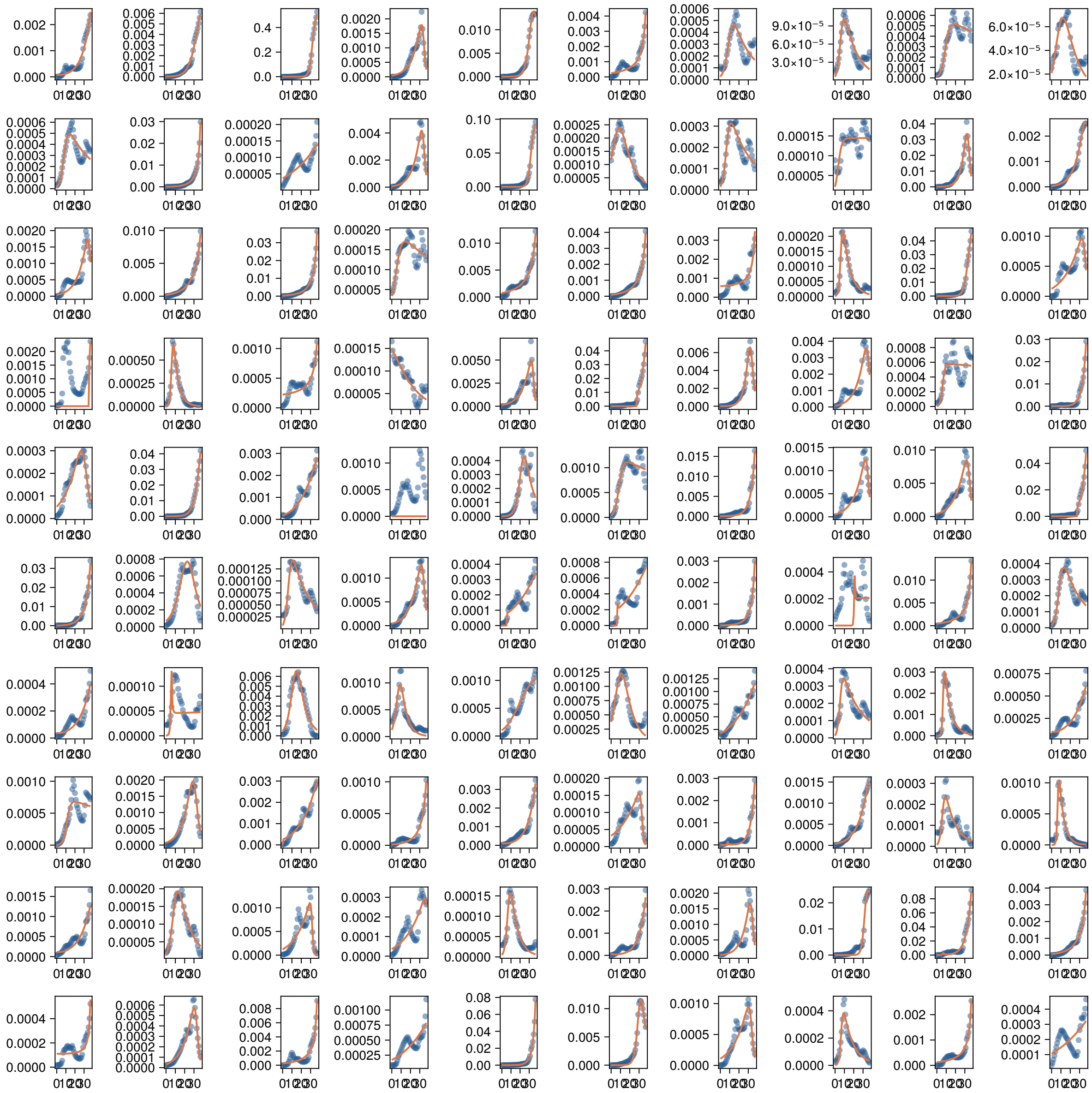

Figure S11: **Emergent TPCs of effective intraspecific interactions ($\alpha_{ii}$) with minimal tradeoff.** Plotted for 100 species competing for 50 resources within a community, across a relatively wider temperature range of 0 - 37 °C with $\rho = 0.0$.

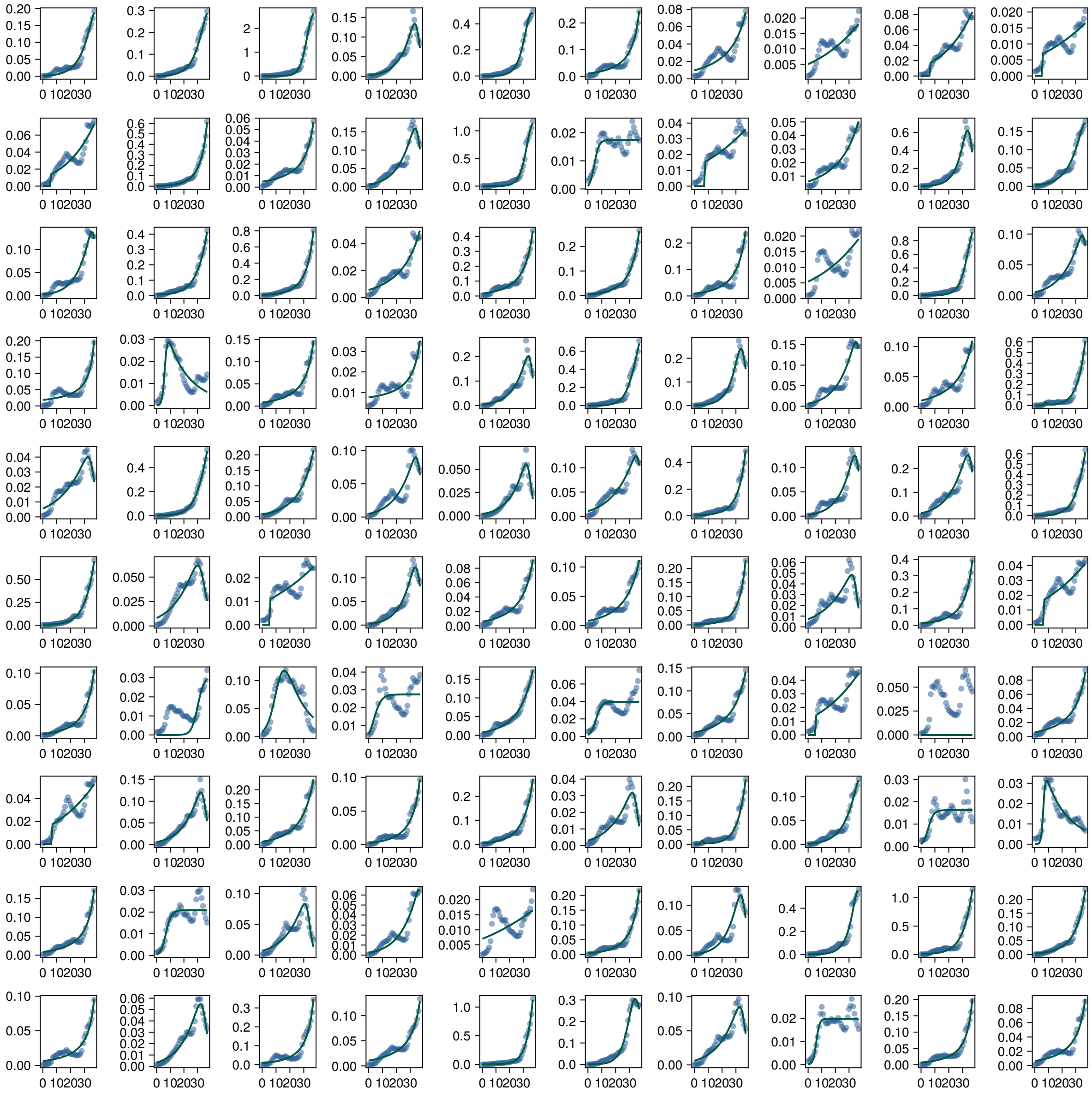

Figure S12: **Emergent TPCs of effective interspecific interactions ($\alpha_{i\neq j}$) with minimal tradeoff.** These TPCs are the impacts on the growth of one species ($i$) by all other species within the community ($\sum_j \alpha_{i\neq j}$), from a 100 species $\times$50 resources community over 0 - 37 °C, for $\rho = 0.0$.

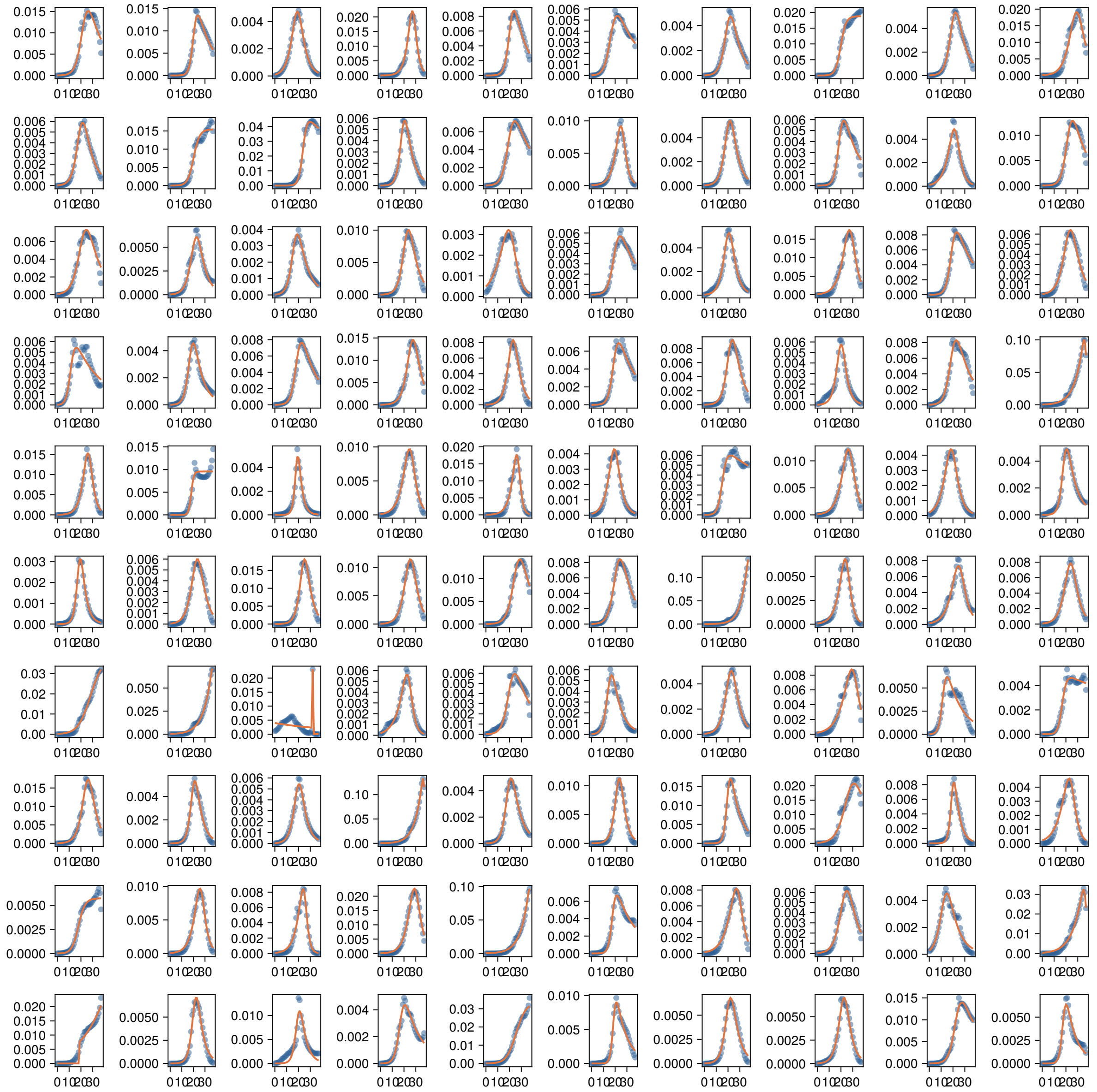

Figure S13: **The emergent temperature performance curves (TPCs) of effective intraspecific interactions ($\alpha_{ii}$).** Plotted for 100 species competing for 50 resources within a community, across a relatively wider temperature range of 0 - 37 °C with $\rho = -1.0$.

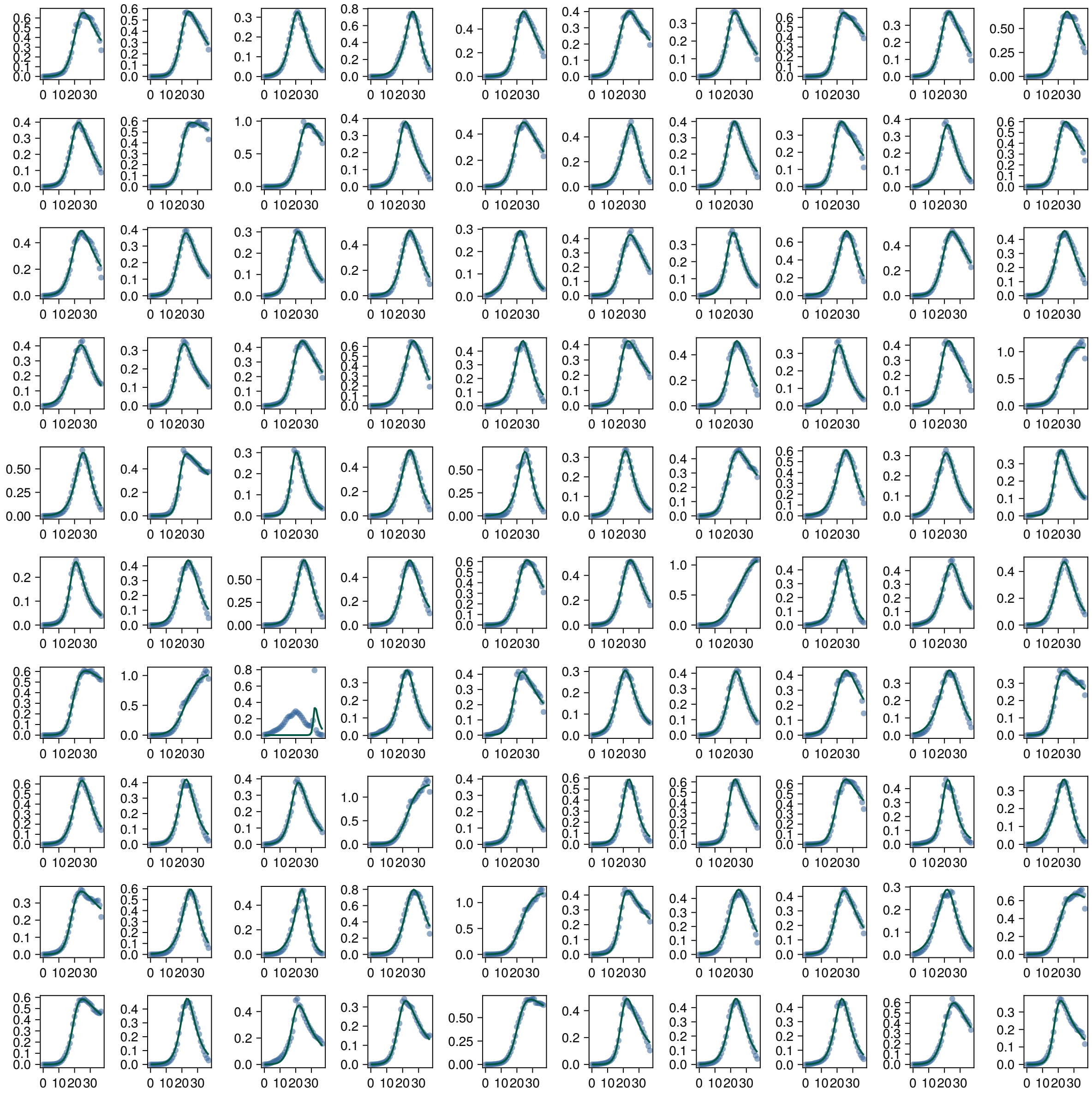

Figure S14: **The emergent temperature performance curves (TPCs) of effective interspecific interactions ($\alpha_{i \neq j}$).** Plotted for 100 species competing for 50 resources within a community, across a relatively wider temperature range of 0 - 37 °C with $\rho = -1.0$. This plot shows the general temperature dependence of the impacts on the growth of one species ($i$) by all other species within the community ($\sum_j \alpha_{i \neq j}$).

### References


Andersen, K. H. & Visser, A. (2023), 'From cell size and first principles to structure and function of unicellular plankton communities', *Progress in Oceanography* **213**, 102995.

Angilletta, M. J., Niewiarowski, P. H. & Navas, C. A. (2002), 'The evolution of thermal physiology in ectotherms', *Journal of thermal Biology* **27**(4), 249–268.

Bag, N., Yap, D. H. X. & Wohland, T. (2014), 'Temperature dependence of diffusion in model and live cell membranes characterized by imaging fluorescence correlation spectroscopy', *Biochimica et Biophysica Acta (BBA)-Biomembranes* **1838**(3), 802–813.

Bellotto, N., Agudo-Canalejo, J., Colin, R., Golestanian, R., Malengo, G. & Sourjik, V. (2022), 'Dependence of diffusion in escherichia coli cytoplasm on protein size, environmental conditions, and cell growth', *Elife* **11**, e82654.

Corkrey, R., Olley, J., Ratkowsky, D., McMeekin, T. & Ross, T. (2012), 'Universality of thermodynamic constants governing biological growth rates', *PLoS One* **7**(2), e32003.

Di Bari, D., Timr, S., Guiral, M., Giudici-Orticoni, M.-T., Seydel, T., Beck, C., Petrillo, C., Derreumaux, P., Melchionna, S., Sterpone, F. et al. (2023), 'Diffusive dynamics of bacterial proteome as a proxy of cell death', *ACS Central Science* **9**(1), 93–102.

Fenchel, T. (1974), 'Intrinsic rate of natural increase: the relationship with body size', *Oecologia* **14**(4), 317–326.

Gibson, B., Wilson, D. J., Feil, E. & Eyre-Walker, A. (2018), 'The distribution of bacterial doubling times in the wild', *Proceedings of the Royal Society B* **285**(1880), 20180789.

Gilchrist, G. W. (1995), 'Specialists and generalists in changing environments. i. fitness landscapes of thermal sensitivity', *The American Naturalist* **146**(2), 252–270.

Huey, R. B. & Hertz, P. E. (1984), 'Is a jack-of-all-temperatures a master of none?', *Evolution* pp. 441–444.

Ketola, T. & Saarinen, K. (2015), 'Experimental evolution in fluctuating environments: tolerance measurements at constant temperatures incorrectly predict the ability to tolerate fluctuating temperatures', *Journal of Evolutionary Biology* **28**(4), 800–806.

Kiørboe, T. (2009), 'A mechanistic approach to plankton ecology', *ASLO Web Lectures* **1**(2), 1–91.

Kontopoulos, D.-G., van Sebille, E., Lange, M., Yvon-Durocher, G., Barraclough, T. G. & Pawar, S. (2020), 'Phytoplankton thermal responses adapt in the absence of hard thermodynamic constraints', *Evolution* **74**(4), 775–790.

Kontopoulos, D., Smith, T. P., Barraclough, T. G. & Pawar, S. (2020), 'Adaptive evolution shapes the present-day distribution of the thermal sensitivity of population growth rate', *PLoS biology* **18**(10), e3000894.

MacArthur, R. (1970), 'Species packing and competitive equilibrium for many species', *Theoretical population biology* **1**(1), 1–11.

Marsland, R., Cui, W., Goldford, J., Sanchez, A., Korolev, K. & Mehta, P. (2019), 'Available energy fluxes drive a transition in the diversity, stability, and functional structure of microbial communities', *PLoS computational biology* **15**(2), e1006793.

Marsland, R., Cui, W. & Mehta, P. (2020), 'A minimal model for microbial biodiversity can reproduce experimentally observed ecological patterns', *Scientific reports* **10**(1), 1–17.

Mills, R. (1973), 'Self-diffusion in normal and heavy water in the range 1-45. deg.', *The Journal of Physical Chemistry* **77**(5), 685–688.

Moloney, C. L. & Field, J. G. (1991), ‘The size-based dynamics of plankton food webs. i. a simulation model of carbon and nitrogen flows’, *Journal of Plankton Research* **13**(5), 1003–1038.

Mongold, J. A., Bennett, A. F. & Lenski, R. E. (1996), ‘Evolutionary adaptation to temperature. iv. adaptation of escherichia coli at a niche boundary’, *Evolution* **50**(1), 35–43.

Padfield, D., O’Sullivan, H. & Pawar, S. (2021), ‘rtpc and nls. multstart: a new pipeline to fit thermal performance curves in r’, *Methods in Ecology and Evolution* **12**(6), 1138–1143.

Pirt, S. J. (1975), *Principles of microbe and cell cultivation.*

Piskulich, Z. A., Mesele, O. O. & Thompson, W. H. (2017), ‘Removing the barrier to the calculation of activation energies: Diffusion coefficients and reorientation times in liquid water’, *The Journal of Chemical Physics* **147**(13).

Schavemaker, P. E., Boersma, A. J. & Poolman, B. (2018), ‘How important is protein diffusion in prokaryotes?’, *Frontiers in molecular biosciences* **5**, 93.

Śmigiel, W. M., Mantovanelli, L., Linnik, D. S., Punter, M., Silberberg, J., Xiang, L., Xu, K. & Poolman, B. (2022), ‘Protein diffusion in escherichia coli cytoplasm scales with the mass of the complexes and is location dependent’, *Science Advances* **8**(31), eabo5387.

Smith, T. P., Thomas, T. J., García-Carreras, B., Sal, S., Yvon-Durocher, G., Bell, T. & Pawar, S. (2019), ‘Community-level respiration of prokaryotic microbes may rise with global warming’, *Nature communications* **10**(1), 5124.

Tang, E. P. & Peters, R. H. (1995), ‘The allometry of algal respiration’, *Journal of Plankton Research* **17**(2), 303–315.

Tilman, D. (1982), *Resource competition and community structure*, number 17, Princeton university press.

Weber, S. C., Spakowitz, A. J. & Theriot, J. A. (2012), ‘Nonthermal atp-dependent fluctuations contribute to the in vivo motion of chromosomal loci’, *Proceedings of the National Academy of Sciences* **109**(19), 7338–7343.
